# Fluorescence recovery after photobleaching reveals different behaviour of tropomyosin isoforms Tpm3.1 and Tpm4.2 in dendritic spines

**DOI:** 10.1101/2025.07.22.666217

**Authors:** T. Tomanić, H. Stefen, E. Parić, A. Chien, T. Fath

**Affiliations:** Dementia Research Centre, Macquarie Medical School, Faculty of Medicine, Health and Human Sciences, Macquarie University, North Ryde, New South Wales Australia; Microscopy Unit, Macquarie Analytical and Fabrication Facility (MAFF), Macquarie University, North Ryde, New South Wales Australia

**Keywords:** actin, cytoskeleton, post-synapse, hippocampal neurons

## Abstract

Actin is the predominant cytoskeletal structure in both the pre- and the post-synaptic compartment of excitatory synapses in the brain, which are formed between the distal part of the axon and the distal site of dendritic spines. Tropomyosin (Tpm) is regarded a master regulator of actin dynamics in mammalian cells. Tpm isoforms, found in neurons are encoded by the *Tpm1*, *Tpm3* and *Tpm4* genes, and have a distinct temporal and spatial distribution of expression. *Tpm3* and *Tpm4* gene products have been found to segregate to the postsynaptic region of central nervous system synapses. Functional differences between Tpm3.1 and Tpm4.2 in neurons have been reported in previous studies. However, these were lacking a detailed analysis of the molecular mobility and dynamics of these two Tpm isoforms in the dendritic compartment. Here, we investigated the kinetic properties of Tpm3.1 and Tpm4.2 via a Fluorescent Recovery After Photobleaching (FRAP) approach and have discovered that Tpm3.1 and Tpm4.2 have distinct kinetic features in dendritic spines. Moreover, we investigated the dynamics of actin in the presence of either Tpm3.1 or Tpm4.2 isoform overexpression, using F-tractin as a reporter of filamentous actin. We have shown that the kinetics of actin turnover is significantly different in response to Tpm3.1 overexpression when compared the actin turnover in response to Tpm4.2 overexpression. Our study further elucidates the roles of Tpm3.1 and Tpm4.2 and provides important conclusions for future studies that are focused on discerning the molecular pathways of Tpm3.1 and Tpm4.2 segregation into different neuronal compartments.

## Introduction

*Tropomyosin (Tpm)* has been regarded as a master regulator of the actin filament function [1]. Over 40 isoforms of Tpm have been identified in mammals, and they are generated by being transcribed and alternatively spliced from four Tpm genes, *Tpm1*, *Tpm2*, *Tpm3* and *Tpm4*. Structurally, Tpm is a helical coiled-coil that polymerises head-to-tail along the actin filament [2]. The N- and C-terminus of the Tpm overlap in the head-to-tail polymer by 11-14 amino acid residues. These termini are the main source of variability between Tpm isoforms, while the rest of the Tpm sequence is highly conserved both between the isoforms and between the species [3]. Tpms have also been classified as high molecular weight (HMW) and low molecular weight (LMW) isoforms, composed out of ∼284 and ∼248 amino acids, respectively [4]. Tpm isoforms in different cells and subcellular compartments each have differing affinities for actin, meaning that they decorate the actin filament in an isoform-specific manner, and this Tpm-actin complex is distinct for various regions within the cell. In this regard, the existence of many Tpm isoforms that are able to diversify the actin filament is considered to be evolutionary advantageous, and the studies so far have suggested that Tpm isoforms are responsible for functions that are non-redundant [5,6].

In central nervous system (CNS) neurons, only three out of the four Tpm genes were found to be active, *Tpm1*, *Tpm3* and *Tpm4.* In neurons, the products of *Tpm1*, Tpm1.10 and Tpm1.12, have shown to be expressed in the pre-synaptic compartment [7–10] while the isoforms with the highest abundance at the post-synapse are Tpm3.1/2 and Tpm4.2 [7,8]. Considering their post-synaptic localisation, and their proposed functions in the literature so far, Tpm3.1 and Tpm4.2 potentially represent a promising target to modulate synaptic functioning in the central nervous system. Although recent studies have addressed the question of the functional roles of Tpm3.1 and Tpm4.2 in Tpm3.1 overexpression [11] and Tpm4.2 knockout [12] mouse models, respectively, the differential roles of Tpm3.1 and Tpm4.2 in regulating the neuronal post-synaptic compartment have not been experimentally addressed.

For the methodological system used in this study (E16.5 cultured mouse hippocampal neurons), it was shown that both *Tpm3* and *Tpm4* gene products segregate to the post-synaptic compartment, while *Tpm1* gene products localize to the pre-synaptic compartment [7]. This, combined with previous research conducted in E18 rat hippocampal neuron cultures, where Tpm4.2 and Tpm1.12 concentrate in the growth cones (with Tpm4.2 more abundant than Tpm1.12) [8], suggests that Tpm4.2 undergoes a developmental shift in its subcellular segregation. Investigations into the distribution of the endogenous Tpm3.1 in mouse primary cultured neurons have shown that Tpm3.1 is abundant in the cell body and primary neurites [13], while it also localises to the growth cones [14]. Exogenously expressed Tpm3.1 was described to be enriched in filopodia and growth cones of mouse primary cortical neurons, where growth cones containing overexpressed Tpm3.1 were particularly enlarged [15]. *Tpm3* gene products have shown to be increased in the brain during embryonic development until birth, after which their levels significantly decrease in the postnatal development [9]. The absence of all LMW *Tpm3* gene products has been demonstrated to be embryonically lethal [16]. Whole brain homogenate experiments showed that levels of both Tpm4.2 and Tpm3.1 decrease during mouse development [17]. At the same time, however, the levels of Tpm4.2 in the PSD fraction progressively increase as mice age, while the levels of Tpm3.1 in the same compartment are significantly decreased in older mice. The combination of these studies points to a necessity for a more detailed understanding of Tpm3.1 and Tpm4.2 functions in the post-synaptic compartment.

Fluorescence Recovery After Photobleaching (FRAP) is a powerful microscopy technique used to analyse the dynamic behaviour of molecules within living cells. It involves selectively photobleaching, or “turning off,” fluorescently labelled molecules in a specific region of interest using a high-intensity laser [18]. By then monitoring the recovery of fluorescence in that region over time, provides valuable insights into molecular mobility and binding interactions [19]. This recovery occurs as unbleached fluorescent molecules move into the bleached area, providing a quantitative measure of the dynamics of the labelled molecules. Photobleaching of dendritic spines in developing mouse primary hippocampal neurons provides some information about the kinetic properties of these two isoforms within the compartment and represents a step closer towards elucidating their roles in neurons. So far, the only published study on Tpms that used FRAP as an experimental tool was Martin et al. (2010) [20]. The research was conducted in NIH3T3 fibroblasts to compare the dynamics of Tpm1.7, Tpm1.9 and Tpm3.1 in stress fibres. The study demonstrated that Tpm3.1 has significantly faster association/dissociation rates with actin filaments when compared to the other two isoforms, indicating that it has lower affinity for associating with actin filament in these cells. All three isoforms had comparable extent of recovery of the mobile fraction. However, FRAP method for the study of Tpms in neuronal cells has not been reported yet. A previous study on rat hippocampal neurons [21] used Fluorescence Decay After Photoactivation (FDAP), a method opposite to FRAP by the means of both biological representation of the acquired signal and curve fitting. The results from this approach combined with other data showed that F-actin in the axon initial segment (AIS) has a lower rate of depolymerisation, and that Tpm3.1 is important for the maintenance of the actin structure within the AIS [21].

Here, we utilize FRAP technique to determine the kinetic properties of Tpm3.1 and Tpm4.2 at the post-synaptic compartment. The findings from this study suggests the presence of functionally distinct Tpm/actin co-polymers within the postsynaptic compartment of cultured hippocampal neurons.

## Materials and Methods

### Primary culture of mouse hippocampal neurons

Cultures of primary hippocampal neurons were prepared as described previously [22]. In short, the neurons were plated on Poly-D-lysine (PDL)-coated 33mm round glass-bottom dishes with 50,000 hippocampal neurons in the centre (plated in 200 µl DMEM, 10% FBS) and 300,000 cortical neurons (plated in 400 µl DMEM, 10% FBS) as a support ring. Two hours after plating, medium was changed to 2 ml complete Neurobasal medium (NBM/B27: Neurobasal, Life Technologies, California, USA; supplemented with 2% B27, Life Technologies, California, USA, and 0.25% GlutaMAX, Invitrogen, Massachusetts, USA) per dish.

### Plasmids and cloning

Adeno-associated viral (AAV) vector used in cloning contains 1.1kb chicken β-actin (CBA) promoter, bovine growth hormone polyadenylation (bGHpA) element and woodchuck hepatitis virus posttranscriptional regulatory element (WPRE), flanked by AAV2 inverted terminal repeats (ITRs) and was previously described in Harasta et al. (2015) [23]. All plasmids were cloned, using NEBuilder Hifi DNA Assembly Master Mix (E2621L, New England Biolabs, Massachusetts, USA) as per manufacturers protocol. For transformations, One Shot Stbl3 chemically competent E. coli (C737303, Thermo Fisher Scientific, Massachusetts, USA) were used and grown in SOC outgrowth medium (B9035, New England Biolabs, Massachusetts, USA). PCRs were performed, using PfuTurbo Cx hotstart DNA polymerase kit (Agilent Technologies, Integrated Sciences). DNA digests were carried out, using restriction enzymes EcoRI-HF (R3101M, New England Biolabs, Massachusetts, USA) and XhoI (R0146M New England Biolabs, Massachusetts, USA), using CutSmart Buffer (B7204, New England Biolabs, Massachusetts, USA). AAV pCAG-mRuby was obtained from Addgene (plasmid #89686). hTpm3.1-mRuby2 was assembled into the AAV vector at the EcoRI restriction site after the amplification of hTpm3.1 sequence from plasmid provided as gift by Peter Gunning (School of Medical Sciences, UNSW, Australia) with the primers 1 and 2, listed in Supplementary Table 1. The amplification of mRuby2 sequence from AAV pCAG-mRuby2 plasmid (Addgene #89686) was performed with primers 3 and 4, listed in Supplementary Table 1. hTpm4.2-mRuby2 was assembled into the AAV vector at the EcoRI restriction site after the amplification of hTpm4.2 sequence from plasmid provided as gift by Peter Gunning (School of Medical Sciences, UNSW, Australia) with primers 5 and 6, listed in Supplementary Table 1. The amplification of mRuby2 sequence from AAV pCAG-mRuby2 plasmid (Addgene #89686) was performed with primers 7 and 8, listed in Table 1. F-tractin-EGFP was assembled into the AAV vector at the EcoRI restriction site after the amplification of F-tractin sequence from pEGFP-C1 F-tractin-EGFP Addgene plasmid #58472 with the primers 9 and 10, listed in Supplementary Table 1. The amplification of EGFP sequence from pEGFP-N1-ACTR3 Addgene plasmid #8462 was performed with primers 11 and 12, listed in Supplementary Table 1. Primers 9, 10, 11, and 12 from Supplementary Table 1. A Kozak sequence was placed in front of the F-tractin sequence, as well as Ser-Gly-Leu-Gly-Ser linker in between the F-tractin and EGFP sequence. For hTpm3.1-IRES-mRuby2 and hTpm4.2-IRES-mRuby2, IRES and mRuby2 sequences were first incorporated into AAV vector at the EcoRI restriction site. IRES sequence was amplified from pCAG GFP IRES Cre plasmid (Addgene #48201), using the primers 13 and 14, listed in Supplementary Table 1. The mRuby2 sequence was amplified from pCAG-mRuby2 plasmid (Addgene #89686) using the primers 15 and 16, listed in Supplementary Table 1. hTpm3.1-IRES-mRuby2 was assembled into AAV_IRES-mRuby2 vector at the XhoI restriction site after the amplification of hTpm3.1 sequence from plasmid provided as gift by Peter Gunning (School of Medical Sciences UNSW, Australia), with the primers 17 and 18, listed in Supplementary Table 1. hTpm4.2-IRES-mRuby2 was assembled into AAV_IRES-mRuby2 vector at the XhoI restriction site after the amplification of hTpm4.2 sequence from plasmid provided as gift by Peter Gunning (School of Medical Sciences, UNSW, Australia), using the primers 19 and 20, listed in Supplementary Table 1.

### Adeno-associated virus (AAV) production and transduction of mouse primary hippocampal neurons

AAV_CAG-mRuby2, AAV_hTpm3.1-mRuby2, AAV_hTpm4.2-mRuby2, AAV_F-tractin-EGFP, AAV_hTpm3.1-IRES-mRuby2 and AAV_hTpm4.2-IRES-mRuby2 were packaged into PHP.eB capsid and the AAV production was performed as previously described [23,24]. After performing qPCR, the following titters were obtained: AAV-PHP.eB_mRuby2 (2.2×10^14^ vp/µL), AAV-PHP.eB_hTpm3.1-mRuby2 (7.6 × 10^13^ vp/µL), AAV-PHP.eB_hTpm4.2-mRuby2 (2.7×10^14^ vp/µL), AAV-PHP.eB_F-tractin-EGFP (1.4×10^14^ vp/µL), AAV-PHP.eB_hTpm3.1-IRES-mRuby2 (9.3×10^13^ vp/µL) and AAV-PHP.eB_hTpm4.2-mRuby2 (2.97×10^14^ vp/µL). For single transductions, the viral stock solution was diluted in NBM and 3.2×10^11^ viral particles were pipetted on 50,000 hippocampal neurons in the centre and 300,000 cortical neurons as support ring, at 3 DIV (AAV-PHP.eB_mRuby2, AAV-PHP.eB_hTpm3.1-mRuby2 or AAV-PHP.eB_hTpm4.2-mRuby2). For double transductions, viral stock solution was diluted in NBM and 1.6×10^8^ viral particles of AAV-PHP.eB_F-tractin-EGFP were mixed with 3.2×10^11^ of either AAV-PHP.eB_mRuby2, AAV-PHP.eB_hTpm3.1-IRES-mRuby2 or AAV-PHP.eB_hTpm4.2-IRES-mRuby2, and pipetted on 50,000 hippocampal neurons in the centre and 300,000 cortical neurons as support ring at 3 DIV.

### Fluorescence recovery after photobleaching

Mouse primary hippocampal neurons were imaged across two days (17-18 DIV), using a ZEISS LSM 880 inverted point-scanning laser confocal microscope and Plan-Apochromat oil objective with 63x magnification (NA 1.4, WD 0.19mm). Lasers used for excitation of EGFP and mRuby2 were Argon (458/488/514nm) and DPCS (561nm) respectively, while the filters used were FITC (Semrock #3540B-zero) and CY3 (Semrock #4040B-zero). The temperature for live imaging was kept at a constant of 37°C and 0.5% CO_2_ supply. The software used for imaging was ZEN Black. After choosing fast smart setup, both 488nm and 561nm lasers were set at 2% strength and 900 gain for F-tractin-EGFP, mRuby2, hTpm3.1-mRuby2 and hTpm4.2-mRuby2 expressing neurons, while 561nm laser was set at 3% strength and 1060 gain for hTpm3.1-IRES-mRuby2 and hTpm4.2-IRES-mRuby2 expressing neurons. The pinhole was set at 1.91 Airy Units (AU), while the zoom was set at 4.0, image size at 680×680 pixels and the speed at 12 (this results in pixel size of 50nm and acquisition speed of 0.561s/frame). Pre-bleaching was set for 10 frames (cycles). No repetitive bleaching was chosen, while the number of iterations was set at 50. The post-bleach acquisition time was set for 214 cycles. The region of 200×200 pixels in the shape of a rectangle was chosen for acquisition and analysis, while the bleaching region was 35×35 pixels in the shape of a circle. The bleaching was performed at 100% laser power (488nm and 561nm laser) and the recovery was measured in the red (561nm) channel for dendritic spines of neurons, transduced with mRuby2, hTpm3.1-mRuby2 and hTpm4.2-mRuby2. For neurons, double transduced with F-tractin-EGFP + mRuby2, F-tractin-EGFP + hTpm3.1-IRES-mRuby2 and F-tractin-EGFP + hTpm4.2-IRES-mRuby2, the recovery was measured in the green (488nm) channel only for those dendritic spines that emerged from neurons positive for mRuby2. Dendritic spines to be bleached were carefully selected to have approximately the same distance from the soma, in those parts of dendrites, where at least one primary branch emerged from the dendritic shaft. More than 40 spines from 4 different glass-bottom dishes in total per experimental group were bleached per each biological replicate, with the total of 3 biological replicates (the total of over 120 spines per one experimental group).

### Imaging

Airy scan imaging of 17-18 DIV mouse primary hippocampal neurons was performed with a confocal laser scanning microscope Zeiss LSM 880 (Carl Zeiss AG, Oberkochen, Germany), equipped with an Airy scan detection unit. To maximise resolution enhancement, a high NA oil immersion Plan-Apochromat 63x/1.46 Oil DIC M27 objective was used. All imaging was performed, using Immersol 518 F immersion media (n_e_ = 1.518, 37°C). Airy scan calibration was done by following the Zeiss manufacturer operation manual before the imaging acquisition started. During the acquisition, an argon laser (488nm) with 0.8% power, and a diode pumped solid state laser (561nm) with 0.8% power were used for excitation. Dual bandpass filter MBS488/561 was applied to separate two fluorescent channels. Bidirectional scanning mode was applied as well, while detector gain and pixel dwell time were adjusted to 880 and 2.06µs, respectively, to avoid phototoxicity and photobleaching effects. Detector digital gain and offset were not applied during the acquisition. To optimise imaging results from the Airy scan, image size was set at 1024×1024 pixels with 1.8 zoom, resulting in a pixel size of 0.07µm (xy lateral resolution). Pinhole size was widely open, as recommended for Airy scan imaging (3.25 AU and 2.94 AU, respectively). Multiple positions and series of z-planes were captured. To further improve the resolution of the acquired images, Airy scan 2D-processing module in Zen Black software was used, with a correction factor of 7.1. All the 2D-processed image stacks from different positions were then projected (with a maximum z-projection) by using ImageJ software and stitched together, using Stitching plugin in ImageJ (v2.1.0, pairwise stitching option).

### Statistical analysis

Significance was determined, using GraphPad prism software (version 9.1.2) by first determining whether the data from each experimental group follows Gaussian distribution with the following tests: Anderson-Darling, D’Agostino & Pearson, Shapiro-Wilk, and Kolgomorov-Smirnov test. After determining that at least one of the experimental groups used for comparisons does not follow Gaussian distribution, the data were statistically analysed with Mann-Whitney U non-parametric unpaired test.

### Data analysis

Videos, acquired in ZEN Black, were opened in ImageJ (v2.1.0) and the fluorescence intensity data were extracted from the bleaching, reference and background region using ROI manager in ImageJ software. A Bleaching region (BL) in the shape of a circle was chosen with the same size and placement on the bleached spine, using ZEISS LSM 880 (35×35 pixels). Reference region (REF) was 15×15 pixels and was placed at the fluorescing area that was not bleached (part of the neuron branch downstream or upstream of the bleached spine). Background region (BG) was the same size as the REF and was placed at the non-fluorescing area (empty space between the neuron branches). Fluorescence intensities were gathered across all 600 imaged frames from all three regions. After subtractions of the BG intensities from REF (REF-BG) and BL (BL-BG) intensities for each frame, normalisation of the data was obtained by dividing the BL-BG by REF-BG values for each frame. To make the recovery curve start from zero, the value from the 11^th^ frame (first post-bleaching frame) was subtracted from [(BL-BG)/(REF-BG)] values in each frame. Pre-bleaching intensity was averaged from the values of the first 10 frames (PREbl), while {[(BL-BG)/(REF-BG)]-11^th^} values were then divided by the PREbl value for each frame. This sets the averaged PREbl values to the 100% fluorescence intensity when compared to the post-bleach recovered fluorescence values. Please note that the 11^th^ frame is not the perfect representation of the actual bleaching moment (zero intensity) but can be used as a very good approximation of the bleaching moment, because the sampling is very dense. This results in normalisation of data that were then fitted to one component (OC) exponential equation:

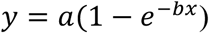

where:

*y* – is fluorescence intensity as a function of time [%];
*a –* is the mobile fraction (the fraction of molecules that can move) [%];
*x* – is the elapsed time [s];
*b* – is a rate constant [s^-1^].

and two component (TC) exponential equation:

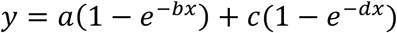

where:

*y* – is fluorescence intensity as a function of time [%];
*a –* is the mobile fraction of slow molecular subpopulation [%];
*x* – is the elapsed time [s];
*b* – is a rate constant of slow molecular subpopulation [s^-1^];
*c* – is the mobile fraction of fast molecular subpopulation [%];
*d* – is the rate constant of fast molecular subpopulation [s^-1^].

Comparison of the goodness of the fit between these two equations is shown in Supplemental Figure 1.

### Programming code

The programming codes used in this article (One-component and Two-component exponential fitting) are uploaded to online repository and can be found at https://osf.io/7eubq/. The goodness of the fit for both of these is explained in the Supplemental Section 1.2.

### One-component (OC) exponential fitting

Plotting and fitting of the data was performed, using the codes, written in MatLab R2020b software. For OC fitting, starting with the section “`Preparation` `of` `the` `data`”, the command “`prepareCurveData`” transforms the data for curve fitting with the fit function. It returns data as columns regardless of the input shapes, during which the number and size of the elements need to match. Data preparation is stretching from line 1 to line 3 in the code. In code section `Set` `up` `fittype` `and` `options`, in line 6, the custom nonlinear fit type was constructed (OC exponential recovery) with additional options, such as designating independent variables for x and dependent variable y (making it dependent of the x value). In line 8, the options of the fit are further defined, such as non-linear least squares fitting method with creating upper and lower limits of the recovery curve. In this regard, `’Lower’, [0, 0]` argument is conditioning the parameters *a* and *b* of the exponential equation to be positive, while `’Upper’, [2, 3]` argument is conditioning parameters *a* and *b* not to go above the values of 2 and 3, respectively. This means that the extent of recovery cannot go beyond 200%, and that the rate constant cannot go beyond 3 [s^-1^]. These were the conditions for OC recovery fitted to the data obtained from the experiment for the recovery of F-tractin. For the OC exponential recovery of mRuby2, Tpm3.1-mRuby2 and Tpm4.2-mRuby2, the lower limit was the same, `’Lower’, [0, 0]`, while the upper limit was `’Upper’, [1, 3]`. This sets the same conditions for the rate constant, while the extent of recovery cannot go beyond 100%. Setting the limitations for OC equation was not necessary, but the reason to include the limiting conditions was, due to the comparisons between OC and TC equations (Supplemental Figure 1), to keep the limitations consistent with the ones necessary for TC equations. Further in this section of the code, in line 10, the display of detailed information about the progress of solving the equation during the run was prevented. Lines 12-13, `Fit` `model` `to` `data`, fit the OC exponential recovery equation to the input data. The commands also save the calculated *a* and *b* parameters as `fitresult`, and goodness of the fit as `gof` variables. Lines 15-30 plot fitted data and display it as a graph. Commands `figure`, `plot`, `legend`, `xlabel`, `ylabel` and `grid` create the new graph window and specify the design of the figure. Other arguments in this section select the font size for the axis labels, legends, numbering, as well as the dot and line size.

### Two-component (TC) exponential fitting

For the TC exponential fitting, the programming code is largely the same as the one for OC exponential fitting (lines 1-3, and 19-36). The changes are apparent in the section that sets the `fittype`, and the section that `fits` `the` `model` `to` `data`. There is an additional definition of variables (lines 5-7) that `Set` `parameter` `c` for each sampling rate condition. This value is set to be the value that **y** takes when **x = 3** seconds. Because **x** is plotted as number of frames, 0.214s/frame sampling rate reaches 3 seconds after 14 frames. Variable **x_z** always has the same realistic value for every investigated dendritic spine, which is the value of 3 seconds. However, every spine will have a different value of **y** that corresponds to **x_z**, meaning that each spine has its own unique amount of recovered fluorescence intensity after 3 seconds of post-bleach recovery, which is represented in variable **z**. This becomes important in the selection of fit-type and conditions of the TC exponential fitting (lines 9-14), where one molecular subpopulation is referred to as the “slower”, while the other as the “faster” fraction. After choosing the same non-linear least square fit-type method as for OC fitting and defining **x** as an independent variable while the **y** variable is dependent on **x**, the upper and lower fitting conditions were set for each parameter. In line 12, `’Lower’, [0, 0, z, 0]` and `’Upper’, [2, 1, z, 3]` conditions set the parameter *a* between the values 0 and 2, parameter *b* between the values 0 and 1, parameter *d* between the values 0 and 3, while the parameter *c* always equals the value of *z*. This means that the extent of recovery of slower fraction (*a*) can reach up to 200%, the rate constant of slower fraction (*b*) can have the value between 0 and 1 [s^-1^], the rate constant of faster fraction (*d*) can reach the value of up to 3 [s^-1^], while the extent of recovery of faster fraction (*c*) will always have the value of recovered fluorescence that is reached after 3 seconds of recovery. This condition makes the TC exponential recovery fitting very robust, and it was necessary to implement it because, more loose conditions give different outcomes for the values of *a*, *b*, *c*, and *d* after each repetitive code run. This means that, when tested empirically, MatLab gave a lot of possible solutions for the values of parameters *a*, *b*, *c*, and *d* in the absence of the fitting limitations, after multiple runs of the code on the same data set. However, if many solutions were given as a result of fitting, one would not know which result to choose from, and random choice would lead to improper comparisons between the experimental groups. This is why conditional arguments need to be tested empirically for every data set and through many repetitions, until the limitations are robust enough so that each repetition starts giving the same result. In this case, the same fitting conditions were applied to 10 data sets from each experimental group for at least 50 code-runs per data set. Once all the code-runs per data set gave the same result for *a*, *b* and *d* (up to the third decimal), the fitting conditions were used for the rest of the data obtained from the experiment. The condition for parameter *c* is, therefore, personalised for every dendritic spine and this makes the comparison between the experimental groups feasible. In other words, fluorophores within each dendritic spine consist of two distinct molecular subpopulations: one that has lower rate constant (up to 1 [s^-1^]) but can recover up to 200% (slower subpopulation), and the other that has three times higher rate constant (up to 3 [s^-1^]), but has a fixed extent of recovery (the value in % that is reached after 3 seconds of recovery). This other subpopulation is referred to as faster subpopulation. Both slower and faster molecular subpopulations recover simultaneously and add up to form an exponential recovery profile that is unique for each dendritic spine. Just like in the case for OC equations, extent of recovery of the raw data in TC equations was observed to be well over 100% in the case of F-tractin-EGFP, which is why the upper limit for parameter *a* was set as the value of 2. The upper limit for parameter *a* in the case of Tpm3.1-mRuby2 and Tpm4.2-mRuby2 was set as the value of 1. This is the only MatLab code change between the experiments. Line 17 of the code fits model to data the same way as for the OC exponential fitting.

## Results

### Tropomyosin molecules display two distinct molecular subpopulations during recovery at the dendritic spine

Primary mouse hippocampal neurons were transduced with Tpm3.1-mRuby2, Tpm4.2-mRuby2 and mRuby2 AAVs at 3 DIV. The dendritic spines were bleached with 100% laser power, and the recovery of fluorophores was imaged at 17-18 DIV (Figure 1). The region of the neuron selected for analysis was the point of the emerging primary dendritic branch (Figure 1, upper left and right panels). The imaging took place upon zooming-in to the chosen dendritic spine, and monitoring the recovery of the fluorophores after bleaching, for the total of 120 seconds (Figure 1, bottom panels). Only those dendritic spines that had comparable fluorescence intensity and surface area were chosen for bleaching (Supplemental Figure S3). Because the recovery behaviour of mRuby2, Tpm3.1-mRuby2 and Tpm4.2-mRuby2 can be better explained with the existence of two molecular subpopulations (as mentioned in section 2.8.2., as well as Supplemental Section 1.2., Supplemental Figure S1), the data gathered from these experiments were fitted to TC equations. In Figure 2H, after fitting all the raw data and putting the results on the same graph, it can be observed that mRuby2 has the fastest recovery and the greatest extent of recovery when compared to Tpm3.1-mRuby and Tpm4.2-mRuby2. Tpm3.1-mRuby and Tpm4.2-mRuby were similar, with the Tpm3.1-mRuby2 recovery slightly slower and with the lower extent of recovery than Tpm4.2-mRuby2.

**Fig 1.**
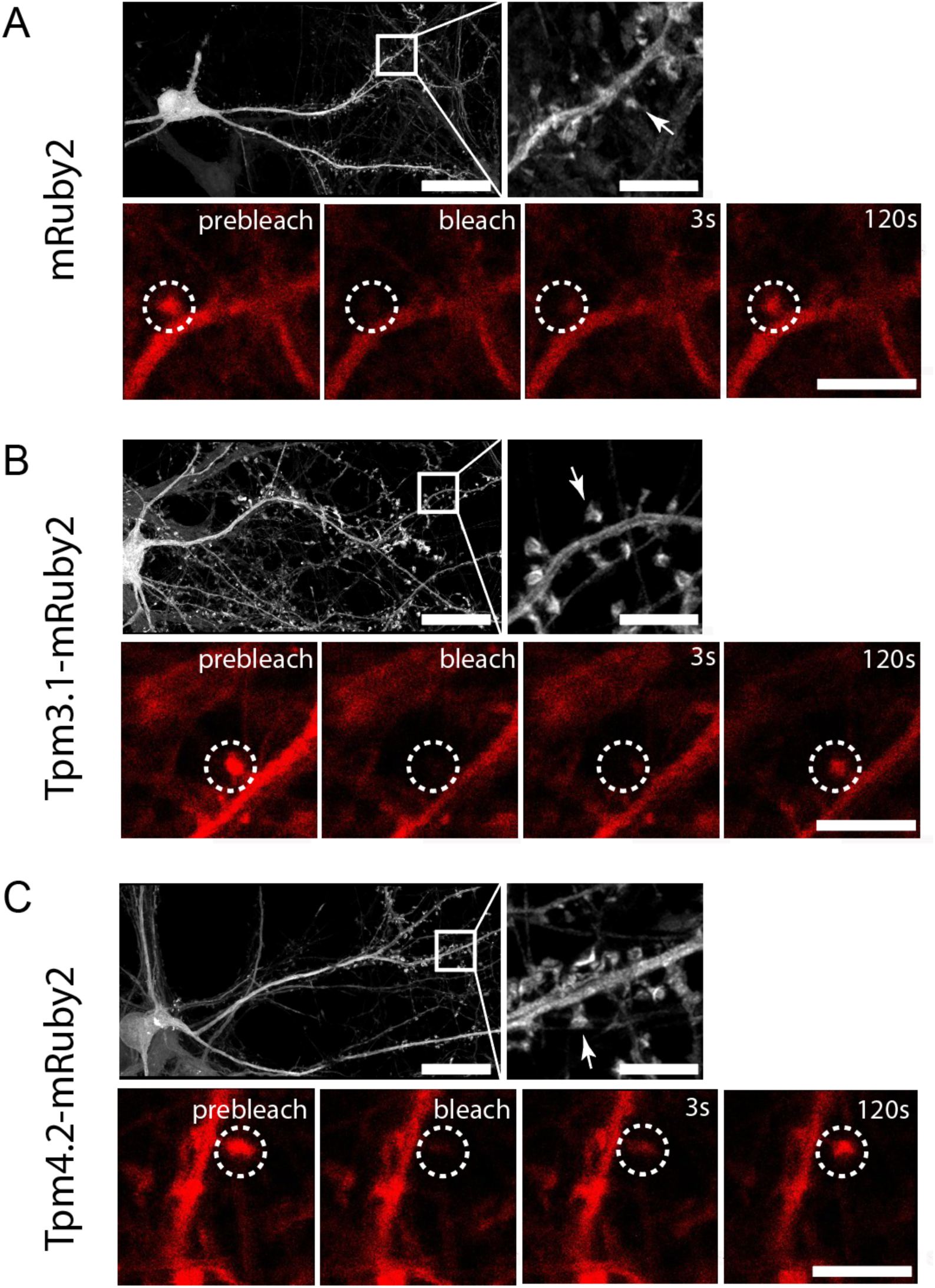
Airy scan (upper panels) and time-lapse images of dendritic spines (bottom panels) of live primary mouse hippocampal neurons transduced with mRuby2, Tpm3.1-mRuby2 or Tpm4.2-mRuby2. Upper left panels: Airy scan image of a neuron with the representative marked area selected for analysis; Upper right panels: zoom-in of the marked area of a neuron, potential spines to be bleached are highlighted with arrows; Bottom panels: Time-lapse images of a bleached dendritic spine highlighted with a circle, with displayed prebleached, bleached, 3s recovery and 120s recovery time points. (A) Neuron transduced with mRuby2. (B) Neuron transduced with Tpm3.1-mRuby2. (C) Neuron transduced with Tpm4.2-mRuby2. Scale bars: Upper left panels = 25μm. Upper right and bottom panels = 5μm

**Fig 2.**
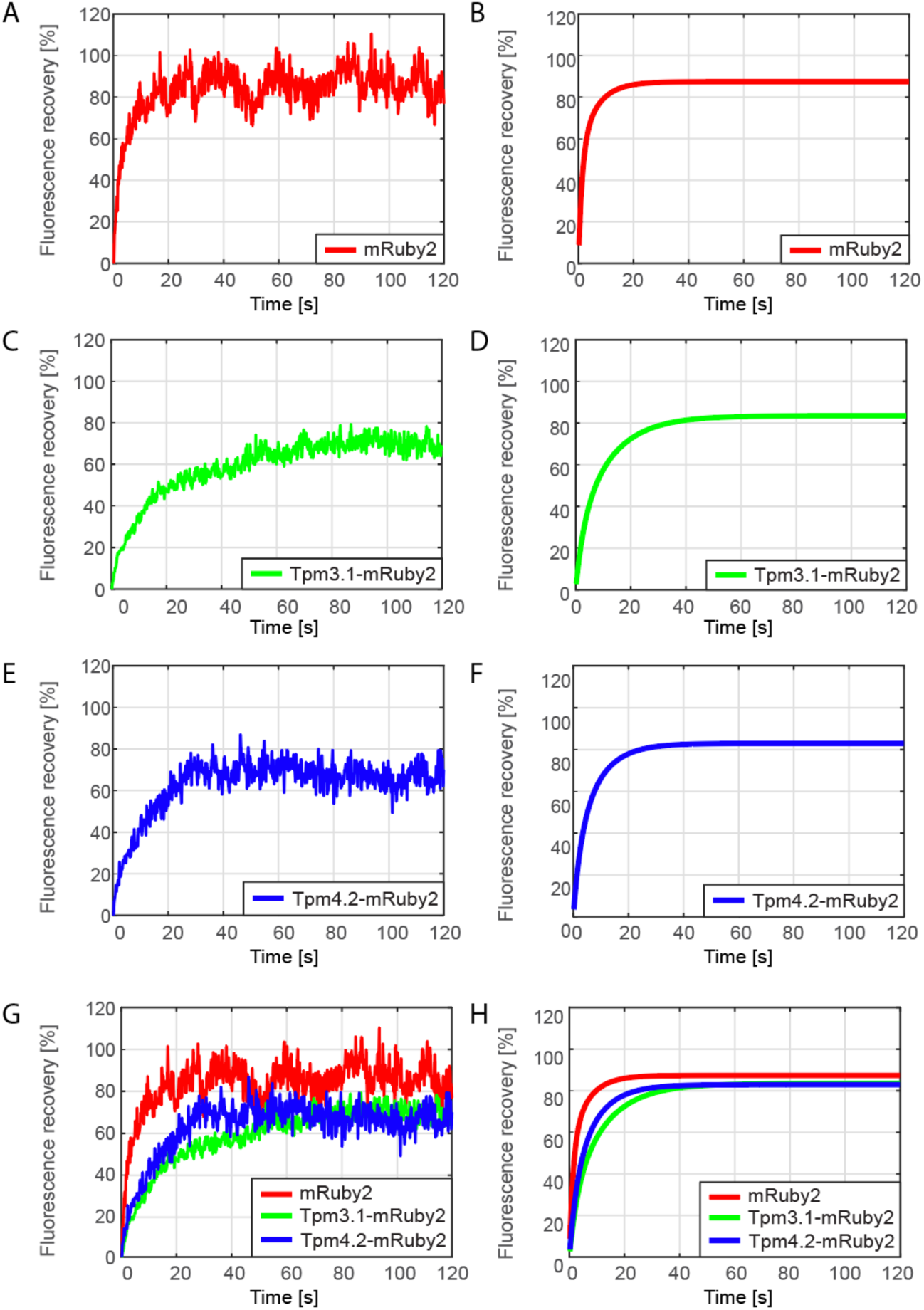
Raw data and Matlab curve-fitted data of bleached dendritic spines of mouse primary hippocampal neurons transduced with mRuby2, Tpm3.1-mRuby2 or Tpm4.2-mRuby2. Red `–` mRuby2 control spines. Green `–` Tpm3.1-mRuby2 spines. Blue `–` Tpm4.2-mRuby2 spines. The data is plotted as the extent of fluorescence recovery (given in [%]), against the time (in seconds) at an experimental sampling rate of 0.214s/frame. (A) Raw data of one representative mRuby2 bleached spine; (B) The TC fitted curves as an average from all of the bleached spines within mRuby2 group; (C) Raw data of one representative Tpm3.1-mRuby2 bleached spine; (D) The TC fitted curves as an average from all of the bleached spines within Tpm3.1-mRuby2 group; (E) Raw data of one representative Tpm4.2-mRuby2 bleached spine; (F) The TC fitted curves as an average from all of the bleached spines within Tpm4.2-mRuby2 group; (G) Raw data of all three representative bleached spines per experimental group shown together; (H) TC fitted curves of each experimental group compared on the same graph. mRuby2 average parameter values: a = 34.53[%], b = 0.03421[s^-1^], c = 52.81[%], d = 0.1568 [s^-1^]. Tpm3.1-mRuby2 average parameter values: a = 57.43 [%], b = 0.01769 [s^-1^], c = 26.06 [%], d = 0.07517 [s^-1^], Tpm4.2-mRuby2 parameter values: a = 52.48 [%], b = 0.02524 [s^-1^], c = 30.43 [%], d = 0.08098 [s^-1^]

As mentioned in section 2.8.2, there are four parameters for TC equations - *a*, *b*, *c*, and *d*: *y* = *a*(1 `−` *e*^−*bx*^) + *c*(1 `−` *e*^−*dx*^). The two molecular subpopulations of fluorophores are recovering simultaneously and are adding up to make one single exponential recovery curve. One of those subpopulations is referred to as the “slower” and represents the fluorophores recovering at the slower rate: *a*(1 `−` *e*^−*bx*^). The other subpopulation is referred to as the “faster” and it represents the fluorophores recovering at the faster rate: *c*(1 `−` *e*^−*dx*^). After 120 seconds of recovery, the slower subpopulation of Tpm3.1-mRuby2 and Tpm4.2-mRuby2 recovered to a significantly higher extent than the control group mRuby2 (Figure 3A). Slower subpopulation of both control mRuby2 and Tpm4.2-mRuby2 recovered at a significantly greater speed than Tpm3.1-mRuby2 (Figure 3B). The faster subpopulation of Tpm4.2-mRuby2 recovered to a significantly greater extent than Tpm3.1-mRuby2, while the amount of recovery for faster subpopulation of control mRuby2 was significantly higher than that of both Tpm3.1-mRuby2 and Tpm4.2-mRuby2 (Figure 3C). Lastly, the speed of recovery for faster subpopulation of Tpm3.1-mRuby2 and Tpm4.2-mRuby2 was significantly lower than the speed of recovery for faster subpopulation of control mRuby2 (Figure 3D).

**Fig 3.**
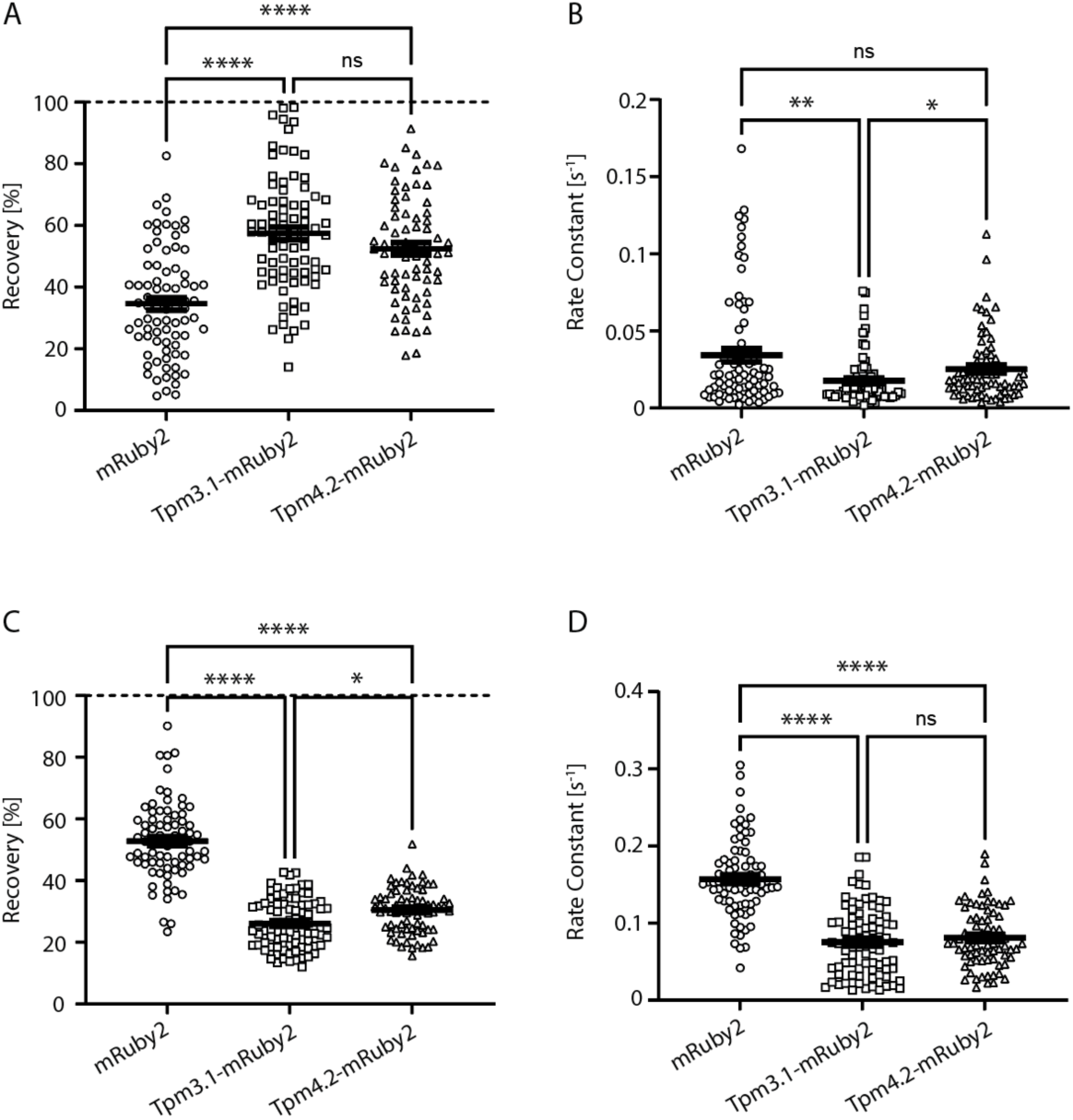
Comparison of TC fitting parameters after 120s of recovery for dendritic spines of primary mouse hippocampal neurons transduced with mRuby2, Tpm3.1-mRuby2 or Tpm4.2-mRuby2. (A) Parameter a `–` extent of recovery for slow subpopulation. (B) Parameter b `–` rate constant for slow subpopulation. (C) Parameter c `–` extent of recovery for fast subpopulation. (D) Parameter d `–` rate constant for fast subpopulation. Significance was determined with Mann-Whitney U non-parametric test: *p<0.05, **p<0.01, ***p<0.001 ****p<0.0001

Please note that the extent of recovery for faster subpopulation is always the same, no matter how long the spines are allowed to recover. This result exists due to the robustness of the programming code used for TC equation fitting (see section 2.8.2).

Overall, the fastest recovery and greatest extent of recovery of mRuby2 when compared to Tpm3.1-mRuby2 and Tpm4.2-mRuby2 is dissected in this section, and it complements Figure 2. As mentioned, mRuby2 is the cytosolic fluorophore and its binding to nearby proteins in the subcellular compartment is not expected, which is confirmed by its fast speed and great extent of recovery in Figure 3. In the case of Tpm3.1-mRuby2, however, its lowest extent of recovery (especially for the slower fraction, Figure 3A) and the lowest speed of recovery (especially for the faster fraction, Figure 3C) can be explained by Tpm3.1 isoform having more binding partners at the dendritic spine than Tpm4.2. These results also complement the representative raw data in Figure 2.

### The recovery of F-tractin molecules with or without the presence of Tropomyosin at the dendritic spine can be explained by the existence of two molecular subpopulations

Similarly to the former experiment (Section 3.1), primary mouse hippocampal neurons were transduced with F-tractin + mRuby2, F-tractin + Tpm3.1-IRES-mRuby2 and F-tractin + Tpm4.2-IRES-mRuby2 AAVs at 3 DIV. The dendritic spines were bleached with 100% laser power, and the recovery of fluorophores was imaged at 17-18 DIV. The region of the neuron selected for analysis was the point of the emerging primary dendritic branch (Figure 4A, 4B, 4C bigger panels). The imaging took place upon zooming-in to the chosen dendritic spine, and monitoring the recovery of the fluorophores after bleaching, for the total of 120 seconds (Figure 4A, 4B, 4C smaller panels, Figure 5). Only those dendritic spines that had comparable fluorescence intensity and surface area were chosen for bleaching (Supplemental Figure S4). As mentioned in the Methods section 2.8. and explained in the Supplementary Section 1.2., the recovery of F-tractin-EGFP molecules can be best fitted to either the OC or TC exponential equations (Supplemental Figure S2). The mean value of adjR2 is trending towards TC exponential fitting, therefore, the recovery of F-tractin + mRuby2, F-tractin + Tpm3.1-mRuby2 and F-tractin + Tpm4.2-mRuby2 was fitted to TC equations. In Figure 6, after fitting all the raw data and putting the results on the same graph (bottom panel on the far right), it can be observed that F-tractin in the presence of mRuby2 has the speed of recovery very similar to that of F-tractin in the presence Tpm3.1, while the F-tractin in the presence of Tpm4.2 has the fastest recovery. The greatest extent of recovery was that of F-tractin in the presence of Tpm4.2, while the F-tractin in the presence of Tpm3.1 showed the lowest extent of recovery. F-tractin in the presence of either Tpm3.1 or mRuby had similar extents of recovery.

**Fig 4.**
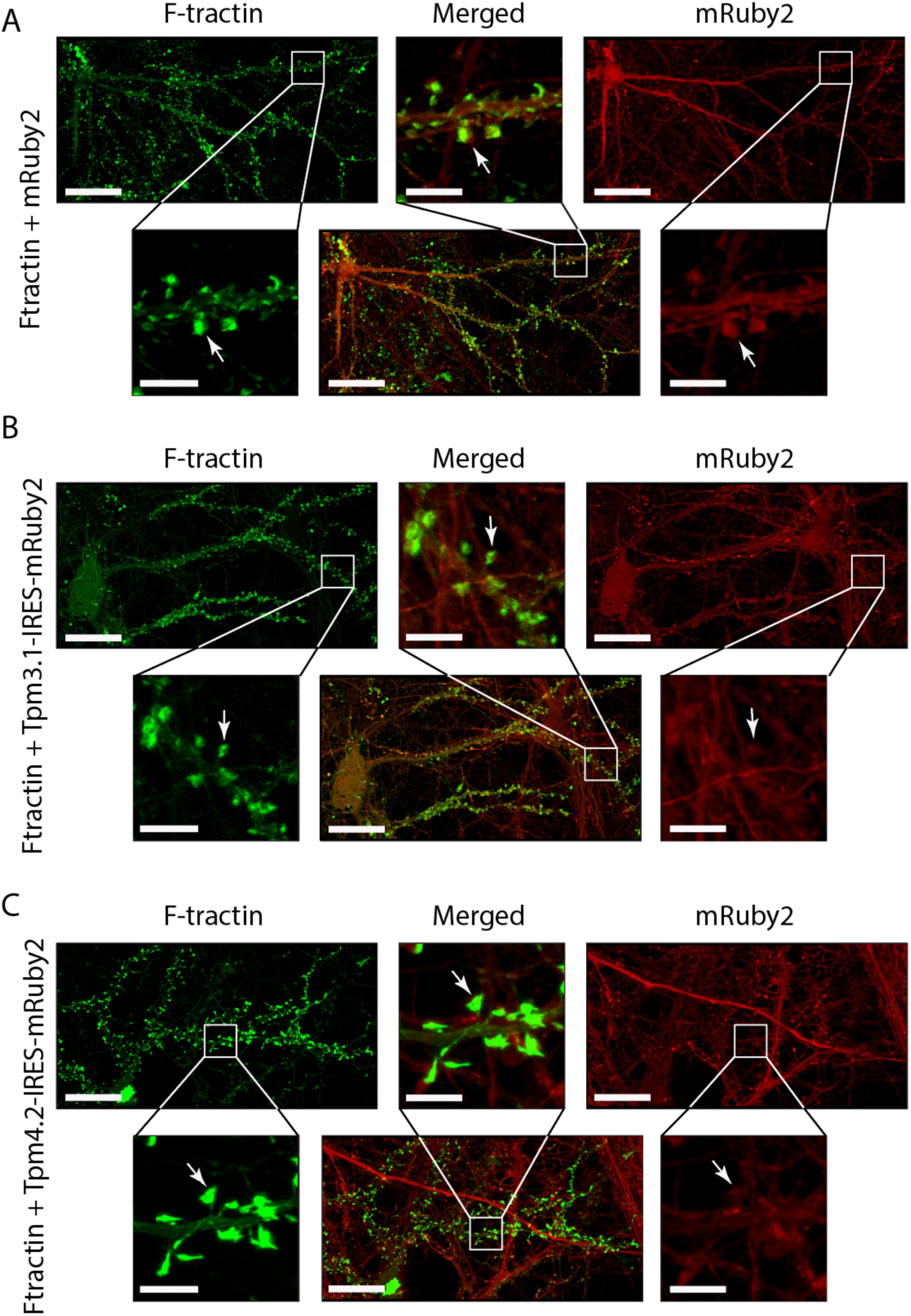
Airy scan images of dendritic spines of live primary mouse hippocampal neurons double transduced with F-tractin-EGFP + mRuby2, F-tractin-EGFP + Tpm3.1-IRES-mRuby2 or F-tractin-EGFP + Tpm4.2-IRES-mRuby2. Upper left panels: Airy scan image of a neuron in green (488nm) channel; Upper right panels: Airy scan image of a neuron in red (561nm) channel; Bottom middle panels: Airy scan image of a neuron in merged green (488nm) and red (561nm) channels; Bottom left panels: zoom-in of the area of a neuron for the dendritic spines to be selected in green (488nm) channel, while spine to be bleached is highlighted with arrows; Bottom left panels: zoom-in of the area of a neuron for the dendritic spines to be selected in red (561nm) channel, while spine to be bleached is highlighted with arrows; Upper middle panels: zoom-in of the area of a neuron for the dendritic spines to be selected in merged green (488nm) and red (561nm) channels, while spines to be bleached are highlighted with arrows. (A) Neurons double transduced with F-tractin-EGFP + mRuby2. (B) Neurons double transduced with F-tractin-EGFP + Tpm3.1-IRESmRuby2. (C) Neurons double transduced with F-tractin-EGFP + Tpm4.2-IRES-mRuby2. Scale bars: Upper left, upper right and bottom middle panels = 25μm. Bottom left, bottom right and upper middle panels = 5μm

**Fig 5.**
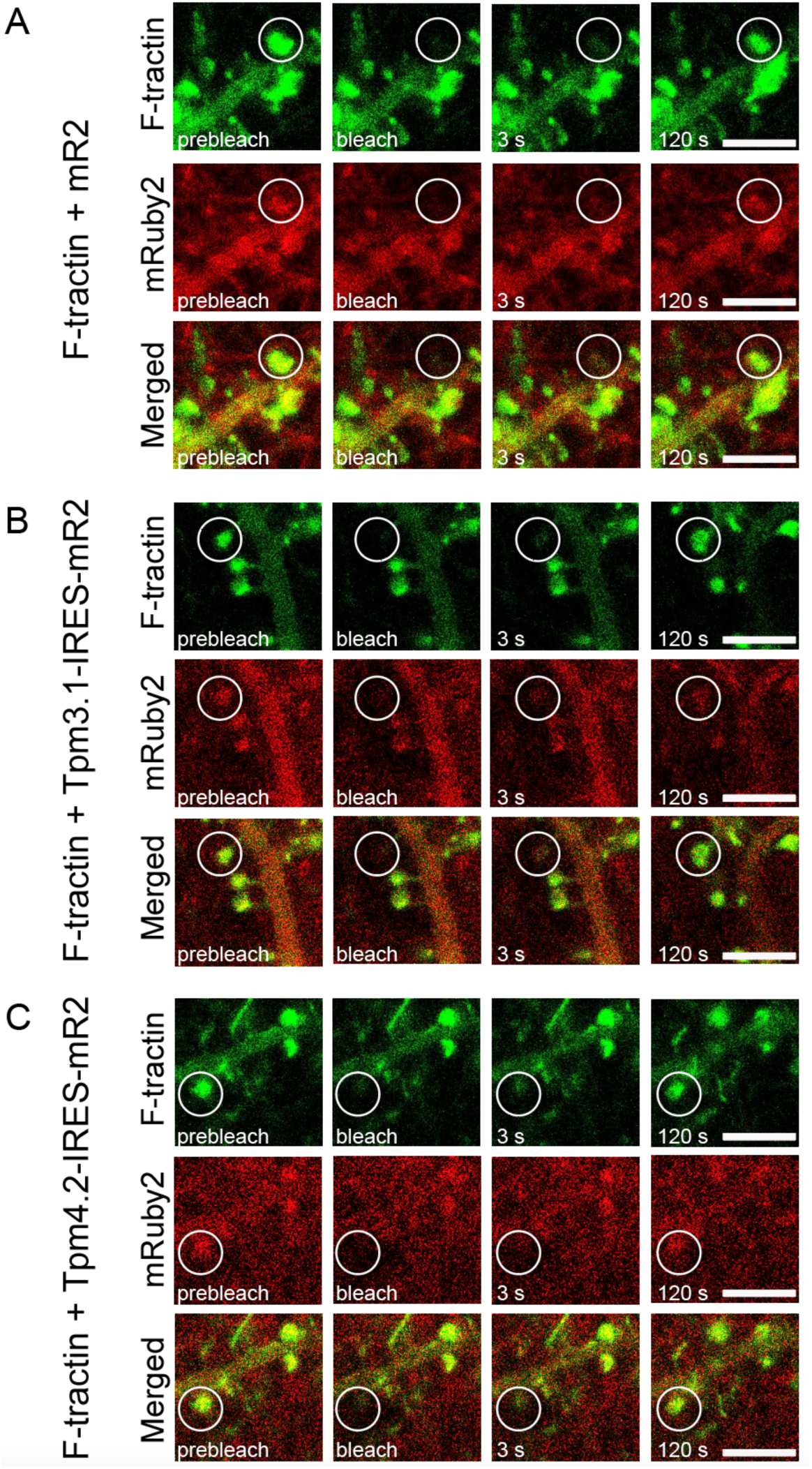
Time-lapse images of bleached dendritic spines of primary mouse hippocampal neurons double transduced with F-tractin-EGFP + mRuby2, F-tractin-EGFP + Tpm3.1-IRES-mRuby2 or F-tractin-EGFP + Tpm4.2-IRES-mRuby2. Bleached dendritic spine is highlighted with a circle, with displayed prebleached, bleached, 3s recovery and 120s recovery time points. Upper panels: bleached dendritic spine shown in green (488nm) channel; Middle panels: bleached dendritic spine shown in red (561nm) channel; Bottom panels: bleached dendritic spine shown in merged green (488nm) and red (561nm) channel. (A) Neurons double transduced with F-tractin-EGFP + mRuby2. (B) Neurons double transduced with F-tractin-EGFP + Tpm3.1-IRES-mRuby2. (C) Neurons double transduced with F-tractin-EGFP + Tpm4.2-IRESmRuby2. Scale bars = 5μm

**Fig 6.**
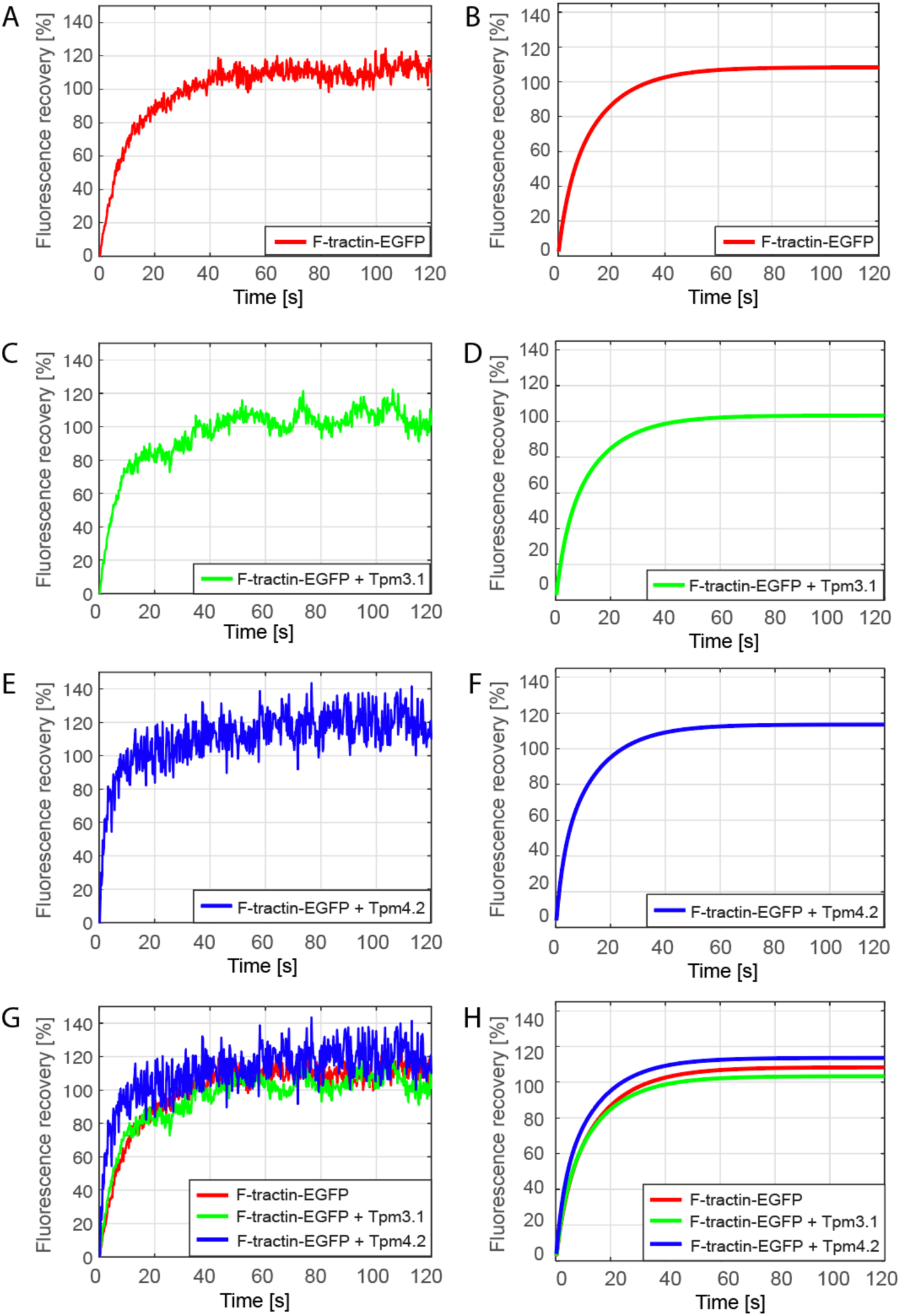
Raw data and Matlab curve-fitted data of bleached dendritic spines of mouse primary hippocampal neurons double transduced with F-tractin-EGFP + mRuby2, F-tractin-EGFP + Tpm3.1-IRES-mRuby2 or F-tractin-EGFP + Tpm4.2-IRES-mRuby2. Red `–`F-tractin-EGFP + mRuby2 control spines. Green `–` F-tractin-EGFP + Tpm3.1-IRES-mRuby2 spines. Blue `–` F-tractin_EGFP + Tpm4.2-IRES-mRuby2 spines. The data is plotted as the extent of fluorescence recovery (given in [%]), against the time (in seconds) at an experimental sampling rate of 0.214s/frame. (A) Raw data of one representative F-tractin-EGFP + mRuby2 bleached spine; (B) The TC fitted curves as an average from all of the bleached spines within F-tractin-EGFP + mRuby2 group; (C) Raw data of one representative F-tractin-EGFP + Tpm3.1-IRES-mRuby2 bleached spine; (D) The TC fitted curves as an average from all of the bleached spines within F-tractin-EGFP + Tpm3.1-IRES-mRuby2 group; (E) Raw data of one representative F-tractin-EGFP + Tpm4.2-IRES-mRuby2 bleached spine; (F) The TC fitted curves as an average from all of the bleached spines within F-tractin-EGFP + Tpm4.2-IRES-mRuby2 group; (G) Raw data of all three representative bleached spines per experimental group shown together; (H) TC fitted curves of each experimental group compared on the same graph. F-tractin-EGFP + mRuby2 average parameter values: a = 79.94 [%], b = 0.0141 [s^-1^], c = 28.38 [%], d = 0.05762 [s^-1^]. F-tractin-EGFP + Tpm3.1-IRES-mRuby2 average parameter values: a = 73.54 [%], b = 0.01481 [s^-1^], c = 29.8 [%], d = 0.06199 [s^-1^], F-tractin-EGFP + Tpm4.2-IRES-mRuby2 parameter values: a = 76.93 [%], b = 0.01529 [s^-1^], c = 36.66 [%], d = 0.07908 [s^-1^]

Equivalently to the former experiment (Section 3.1) the four parameters of TC equations (*a*, *b*, *c* and *d*) were calculated for the 120s recovery time point and compared between the experimental groups. There was no significant difference between the slower subpopulations of F-tractin in the presence of either mRuby2, Tpm3.1 or Tpm4.2 with regards to both the extent and the speed of recovery (Figure 7A, 7B). The proportion of the recovered faster subpopulation of F-tractin was significantly higher in the presence of Tpm4.2 than in the presence of either mRuby2 or Tpm3.1 (Figure 7C). Similarly, the speed of recovery of the faster F-tractin subpopulation in the presence of Tpm4.2 was significantly higher than in the presence of either mRuby2 or Tpm3.1 (Figure 7D).

**Fig 7.**
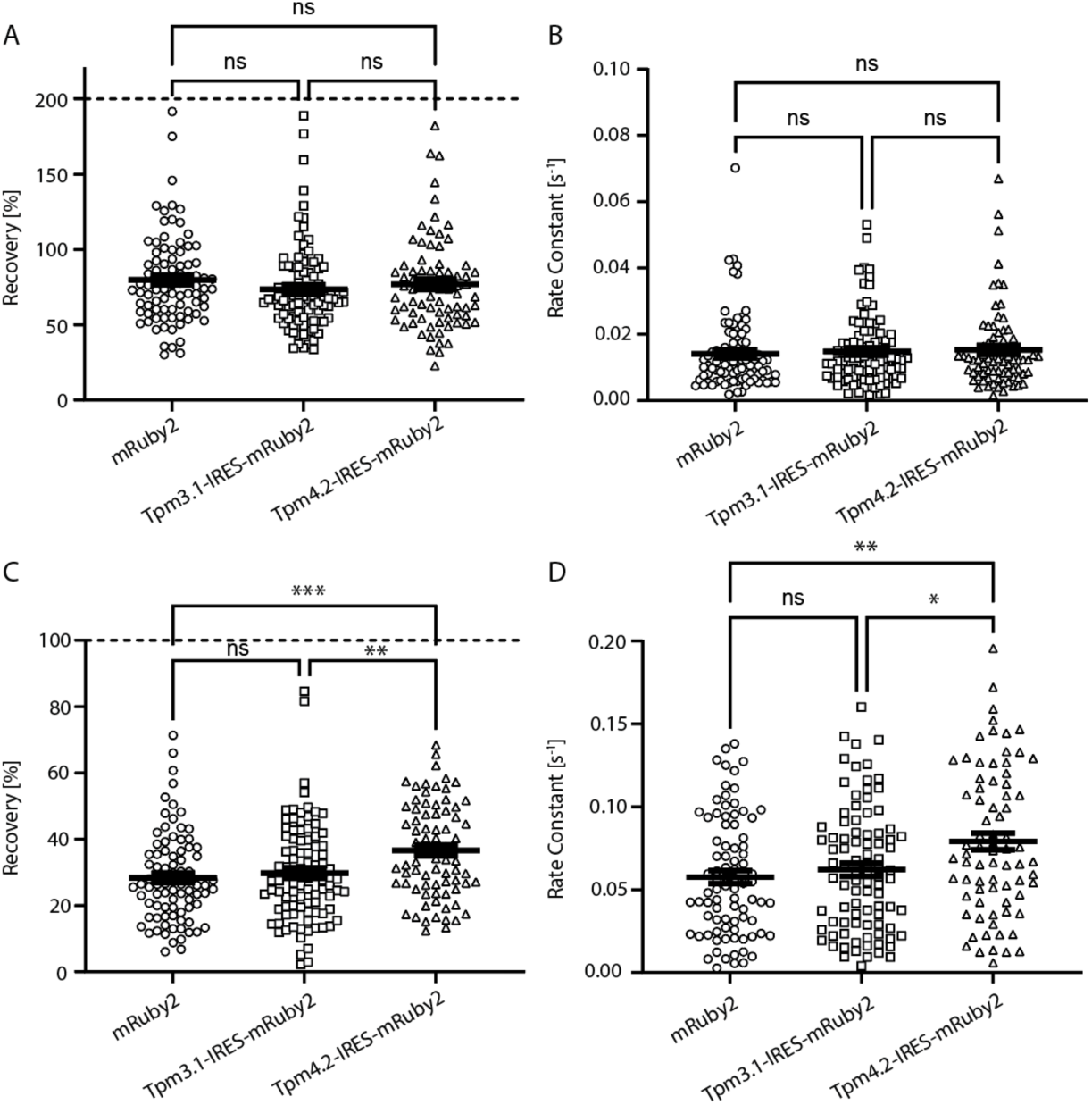
Comparison of TC fitted parameters after 120s of recovery of F-tractin, for dendritic spines of primary mouse hippocampal neurons double transduced with F-tractin-EGFP + mRuby2, F-tractin-EGFP + Tpm3.1-IRES-mRuby2 or F-tractin-EGFP + Tpm4.2-IRES-mRuby2. (A) Parameter a `–` extent of recovery for slow subpopulation. (B) Parameter b `–` rate constant for slow subpopulation. (C) Parameter c `–` extent of recovery for fast subpopulation. (D) Parameter d `–` rate constant for fast subpopulation. Significance was determined with Mann-Whitney U non-parametric test: *p<0.05, ***p<0.001 ****p<0.0001

Please note that the extent of recovery for faster subpopulation is always the same, no matter how long the spines are allowed to recover. This result exists due to the robustness of the programming code used for TC equation fitting (see section 2.8.2).

The quantification of parameters in Figure 7 complements the fitting of raw data displayed in Figure 6. Overall, the extent and speed of recovery for F-tractin were significantly higher in the presence of Tpm4.2 than in the presence of either mRuby2 control or Tpm3.1. This suggests that filamentous actin (which F-tractin has a high affinity to) is less stable, more diffusing and exhibits fewer binding events within the dendritic spine in the presence of Tpm4.2 than in the presence of either Tpm3.1 or mRuby2.

## Discussion

The results from this study show the effective use of FRAP to define isoform-specific properties of individual Tpm isoforms within dendritic spines of cultured neurons. The FRAP approach identified different mobility of the two major Tpm isoforms, Tpm3.1 and Tpm4.2, found in the postsynaptic compartment of excitatory neurons in the brain, and different Tpm3.1 and Tpm4.2-dependent molecular dynamics of the postsynaptic actin cytoskeleton.

### FRAP as tool for assessing the molecular dynamics or functionally related proteins in the postsynaptic compartment

The programming code for two-component exponential recovery fitting was designed to enable proper comparison of kinetic properties between Tpm3.1 and Tpm4.2, and it successfully revealed differences in the dynamic features of these two Tpm isoforms. Two component equations showed either a significantly better fitting than one component equations (Supplemental Figure S1), or a trend towards better fitting than that of one component equations (Supplemental Figure S2). Many FRAP studies on the dendritic spines demonstrated that two (or more) component equations fit better to the raw data. However, the data interpretation of more than one molecular component is challenging, and the studies usually adhere to using only one component to explain the obtained results [25–33,18]. Here, we made the fitting robust to more accurately define the two molecular subpopulations (slow and fast) which allows for suitable comparison of kinetic parameters between the experimental groups. As described in Materials and Methods section 2.8., the fast subpopulation is allowed to recover for 3s, while the slow population recovered for 2 min. Previously, three pools of F-actin were identified in dendritic spines, with the dynamic pool at the tip of the spines having fast turnover (〜40s), a stable pool at the spine base with slow turnover (〜17min), and an enlargement pool in the central spine region whose dynamics change with synaptic activity (2-15min) [34]. This means that, due to the time limit of 2 min, the imaging in our experiments indirectly describes the kinetic properties of only the dynamic pool of actin in dendritic spines, with the recovery of Tpm molecules as master regulators of the actin filament, and the recovery of F-tractin being the reporter of the amount of filamentous actin, present in the spines.

### Differential mobility of Tpm3.1 and Tpm4.2 in dendritic spines

As predicted, we found that the fast population of Tpm3.1 and Tpm4.2 recovers to a significantly lower extent and at a significantly lesser speed than the control mRuby2 (Figure 3C, 3D). This means that after 120 s of recovery, there are more fast-diffusing molecules of the control fluorophore in the dendritic spine than the fast-diffusing molecules of Tpm3.1 and Tpm4.2. The fast subpopulation of the control fluorophore also moves faster, likely because mRuby2 is a freely diffusing molecule, twice smaller than Tpm3.1-mRuby2 and Tpm4.2-mRuby2 molecules, with no binding partners at the dendritic spine. Contrary to mRuby2, the molecules of Tpm3.1 and Tpm4.2 are binding to filamentous actin at the dendritic spine, which reduces their mobility.

Whilst the data strongly suggest that both Tpm3.1. and Tpm4.2 associate with actin filaments in the dendritic spines, the binding of the fast subpopulation of Tpm3.1 is significantly stronger than the binding of the fast population of Tpm4.2 (Figure 3C). The slow population of mRuby2, Tpm3.1 and Tpm4.2 compensates for the observed mobility of their fast subpopulation. The slow subpopulation of Tpm3.1 and Tpm4.2 recovers to a significantly greater extent, at a significantly lower speed than the control mRuby2 (Figure 3A, 3B). This means that after 120 seconds of recovery, there are more slow-diffusing molecules of Tpm3.1 and Tpm4.2 than mRuby2 molecules at the dendritic spine, with the Tpm4.2 molecules diffusing more rapidly than the Tpm3.1 molecules. The binding (association) of Tpm3.1 to its molecular partners is likely starting much sooner than the binding of Tpm4.2, while the dissociation of bleached Tpm4.2 could also be starting sooner than the dissociation of the bleached Tpm3.1. This means that Tpm3.1 is the least capable of contributing to fluorescence recovery from the start of the post-bleaching period. Because the slow and the fast subpopulations add up to constitute one single recovery profile, it can be concluded that most of mRuby2 control molecules recover very fast in the initial stages (fast subpopulation), and a small number of molecules are left to contribute to recovery over the prolonged periods of time (slow subpopulation). The reverse is true for Tpm3.1 and Tpm4.2. It is important to note that the speed of recovery of the Tpm4.2 slow subpopulation is almost the same as the speed of recovery of control fluorophore by 120 seconds of recovery, while Tpm3.1 still recovers at a significantly lower speed by that time (Figure 3B). This suggests that Tpm3.1 continuously binds to filamentous actin at the dendritic spine for the entire duration of the experiment, or that the bleached Tpm3.1 in the bleaching spot hardly dissociates from the interacting partners. At the same time, the Tpm4.2 molecules, have much more rapid association/dissociation events (and are probably replaced by both the exogenous and the endogenous Tpm3.1 at binding sites).

### Differential effect of Tpm3.1 and Tpm4.2 on actin filament properties in dendritic spines

The investigation of properties of filamentous actin using the F-tractin-EGFP reporter also reveals the functional difference between the Tpm3.1 and Tpm4.2 isoforms. After 120 seconds of recovery (Figure 7A), the slow subpopulation of F-tractin under control conditions recovers to an extent that is comparable to the extent of recovery of the slow subpopulation of F-tractin in the presence of either Tpm3.1 or Tpm4.2. The recovery of the F-tractin slow subpopulation also occurs at a similar speed for all three experimental groups (Figure 7B). This means that over prolonged periods of time, the slower subpopulation of F-tractin diffuses and binds to filamentous actin in a similar fashion, regardless of the presence of Tpm3.1 or Tpm4.2. However, the fast subpopulation of F-tractin both under control conditions and in the presence of Tpm3.1, recovers to a significantly lesser extent and at a significantly lower speed when compared to the F-tractin fast subpopulation in the presence of Tpm4.2 (Figure 7C, 7D). This suggests that F-tractin alone is more tightly bound to filamentous actin (F-actin) at the dendritic spine without changing its binding affinity in the presence of Tpm3.1, but it dissociates from F-actin when Tpm4.2 is overexpressed in the cell. The results for the fast subpopulation in Figure 3 show that binding is stronger for Tpm3.1 than for Tpm4.2, but they also do not completely exclude Tpm4.2 from binding events. If F-tractin, Tpm3.1, and Tpm4.2 are all competing for binding with F-actin, the question is why F-tractin is not displaced to the same extent in the presence of Tpm3.1. The answer could be that F-actin is a very dynamic structure and that the event of bleaching, as well as the presence of Tpm3.1 and Tpm4.2, have differing impact on actin remodelling. If bleaching is a stressful event that promotes F-actin polymerisation, it is possible that the presence of Tpm3.1 further stabilizes F-actin structure and polymerisation, creating even more sites for the association of either F-tractin or Tpm3.1. However, the presence of Tpm4.2 does not promote the polymerization of filamentous actin, the number of association sites remains the same as under control conditions, and in the competition for these sites, a certain proportion of F-tractin molecules is displaced from F-actin by Tpm4.2.

### Functionally distinct Tpm3.1/actin and Tpm.4.2/actin co-polymers

The FRAP analyses in this study support the idea that both Tpm3.1 and Tpm4.2 form co-polymers with actin in the postsynaptic compartment with isoform-specific differences in their kinetic parameters. Both Tpm3.1 and Tpm4.2 are not essential for brain development [21,14]. On the molecular level, Tpm3.1 and Tpm4.2 have been shown to have similar physicochemical properties *in vitro*, regarding their influence on F-actin dynamics and their ability to stimulate ATPase activity of NMIIa [35]. However, the *in vitro* characterisation of Tpm3.1 and Tpm4.2 also identified significant differences in how these isoforms regulate F-actin stability and dynamics. Multiwavelength TIRF imaging revealed that Tpm3.1 provides stronger protection from cofilin-induced actin disassembly, compared to Tpm4.2 [36]. Furthermore, myosin motor activity assays, using a NMIIa myosin construct in the presence of labelled F-actin, identified higher sliding velocity of NMIIa along Tpm3.1/actin co-polymers, compared to Tpm4.2/actin co-polymers. The presence of both isoforms in the dendritic spines [7,17], the findings on functional differences of Tpm3.1 and Tpm4.2 *in vitro* [35,36] and the differences in the kinetic parameters of Tpm3.1 and Tpm4.2 in the postsynaptic compartment found in this study, suggest functionally distinct roles of Tpm3.1/actin and Tpm4.2/actin co-polymers at central nervous system synapses. Studies assessing the effect of *Tpm3* and *Tpm4* knockout on neurite growth parameters are consistent with the model of functionally distinct Tpm3.1/actin and Tpm4.2/actin co-polymers in neurons [12,37]. Whilst the knockout of LMW Tpm3 isoforms significantly decreases neurite length and complexity [37], the knockout of Tpm4.2 significantly increases neurite length and complexity of hippocampal neurons [12], cultured for four days.

## Limitations

Our study provides important insights into the differences in the dynamic properties of the two most investigated, neuronally expressed Tpm isoforms in the post-synaptic compartment, Tpm3.1 and Tpm4.2. However, it is pivotal to acknowledge the limitations of our research. Whilst Tpm3.1 is not the only *Tpm3* product at the post-synapse, Tpm3.1/2 show the highest abundance in this compartment when compared to all other *Tpm3* products [17]. In addition, the FRAP analysis was carried out at a single developmental time point. Due to the isoform expression being developmentally regulated, future studies may analyze the differential role of Tpm3 and Tpm4 isoforms in the post-synapse in neurons cultured for different lengths of time.

## Conclusion

The application of FRAP in the present study has been instrumental in resolving distinct dynamic properties of the major post-synaptic tropomyosin isoforms Tpm3.1 and Tpm4.2. By enabling the quantitative measurement of protein mobility in living cells, FRAP allowed us to distinguish key differences in the turnover rates and binding dynamics of Tpm3.1 and Tpm4.2 at dendritic spines. These differential mobility profiles point to isoform-specific roles in the regulation of synaptic architecture or plasticity. A direct comparison of changes in synaptic function after manipulation of Tpm3.1 and Tpm4.2 protein expression are currently lacking. The current findings suggest that Tpm3.1 and Tpm4.2 have distinct roles in postsynaptic organization. However, the precise mechanisms by which they influence receptor trafficking, actin remodelling, or synaptic signalling remain unclear. Exploring the interactions of Tpm3.1 and Tpm4.2 with known postsynaptic scaffolding proteins or signalling complexes will be critical for uncovering pathways that selectively engage one Tpm isoform over the other.

## Acknowledgements

We thank Professor Roland Brandt at the University of Osnabrück for his critical feedback to the data analysis in this study. We would also like to thank the Macquarie University Central Animal Facility personnel for animal husbandry and Prof Yazi Ke and Ms Yijun Lin for AAV production.

## Supplementary Information

### 1 Supplementary Methods

#### 1.1. Table 1 – primers used for cloning

**Table 1.**
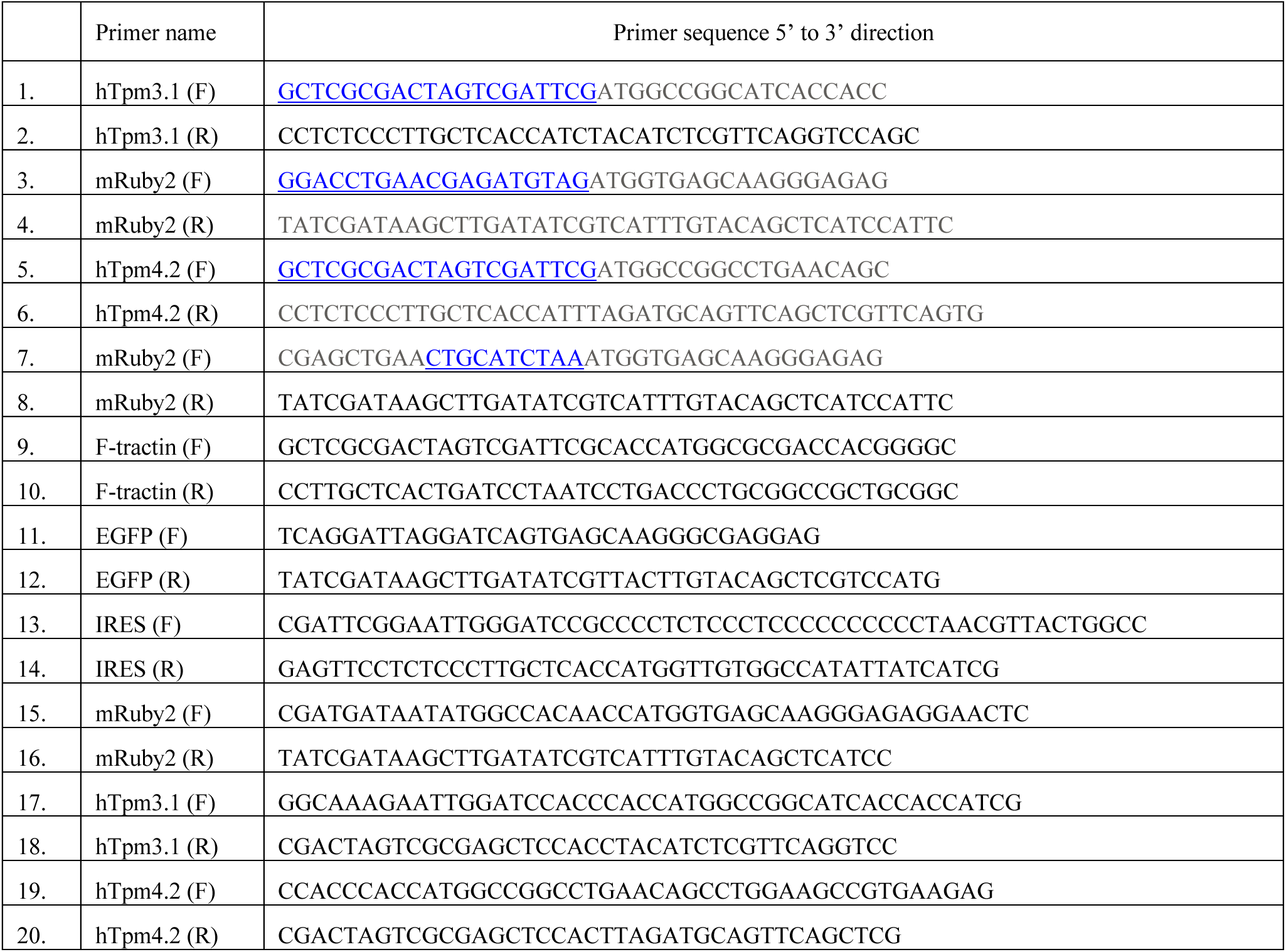
Primer sequences used for cloning of plasmids applied in FRAP experiments. The sequence of the primers is outlined in 5’ to 3’ (left to right) direction. Forward primers put the Kozak sequence upstream of the start codon, while the reverse primers put stop codon downstream of gene sequence. Further explanation is in the main text. F: forward primer. R: reverse primer.

#### 1.2. Goodness of the fit

After fitting the data to the OC and TC recovery exponential equations, the adjusted R2 value (adjR2) was calculated to serve as a parameter of evaluating the goodness of the fit (GOF). The adjR2 values were extracted from gof command given in the programming code. The closer the value of adjR2 gets to the value of 1, the better the fit is. In the case of mRuby2, Tpm3.1-mRuby1 and Tpm4.2-mRuby2, the difference was significant, and it was in favour of the TC fitting equations for all three experimental groups (Fig. S1). This means that the existence of two molecular subpopulations (slower and faster) can explain the recovery of mRuby2, Tpm3.1-mRuby2 and Tpm4.2-mRuby2 molecules after bleaching much better than the existence of only one molecular population. In the case of F-tractin + mRuby2, F-tractin + Tpm3.1-IRES-mRuby2 and F-tractin + Tpm4.2-IRES-mRuby2, there was no significant difference between the adjR2 values when data are fitted to either OC or TC equations for any experimental group (Figure S2). This means that the existence of either one or two molecular subpopulations can equally well explain the recovery behaviour of molecules F-tractin + mRuby2, F-tractin + Tpm3.1-IRES-mRuby2 and F-tractin + Tpm4.2-IRES-mRuby2. However, the overall trend of the fitting still goes in favour of the TC equations for latter experimental groups (the mean value of adjR2 for TC fitting is consistently higher than the one for OC fitting in all three experimental groups – see Supplemetal Figure S2A, S2B, S2CA). Therefore, F-tractin + mRuby2, F-tractin + Tpm3.1-IRES-mRuby2 and F-tractin + Tpm4.2-IRES-mRuby2 were also fitted to TC equations.

**Figure S1.**
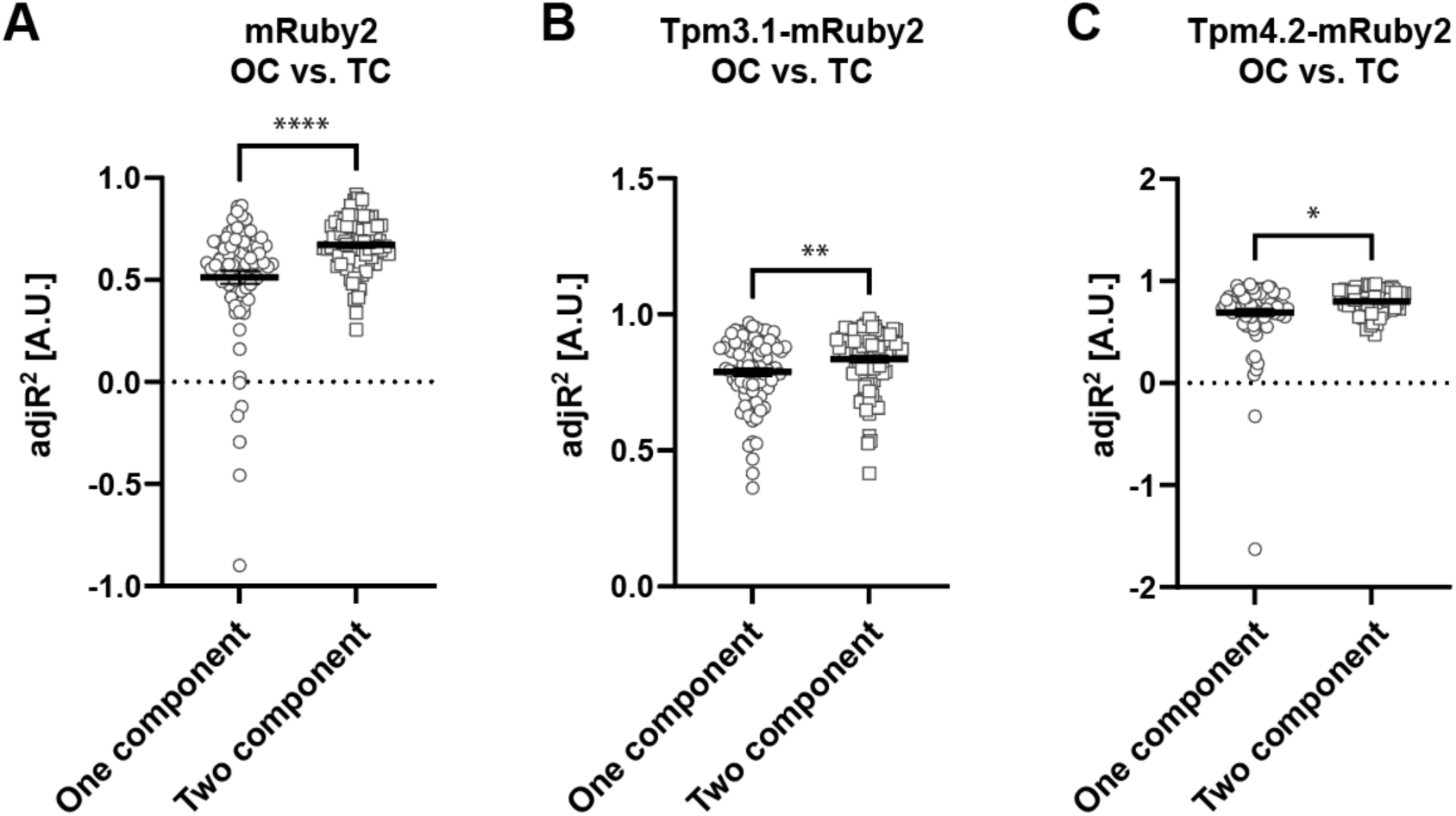
Goodness of the fit (adjR^2^) of OC and TC equations for dendritic spines of primary mouse hippocampal neurons transduced with mRuby2, Tpm3.1-mRuby2 and Tpm4.2-mRuby2 after 120s of recovery. (A) Spines transduced with mRuby2. (B) Spines transduced with Tpm3.1-mRuby2. (C) Spines transduced with Tpm4.2-mRuby2. Significance was determined with Mann-Whitney U non-parametric test: *p<0.05, **p<0.01, ****p<0.0001.

**Figure S2.**
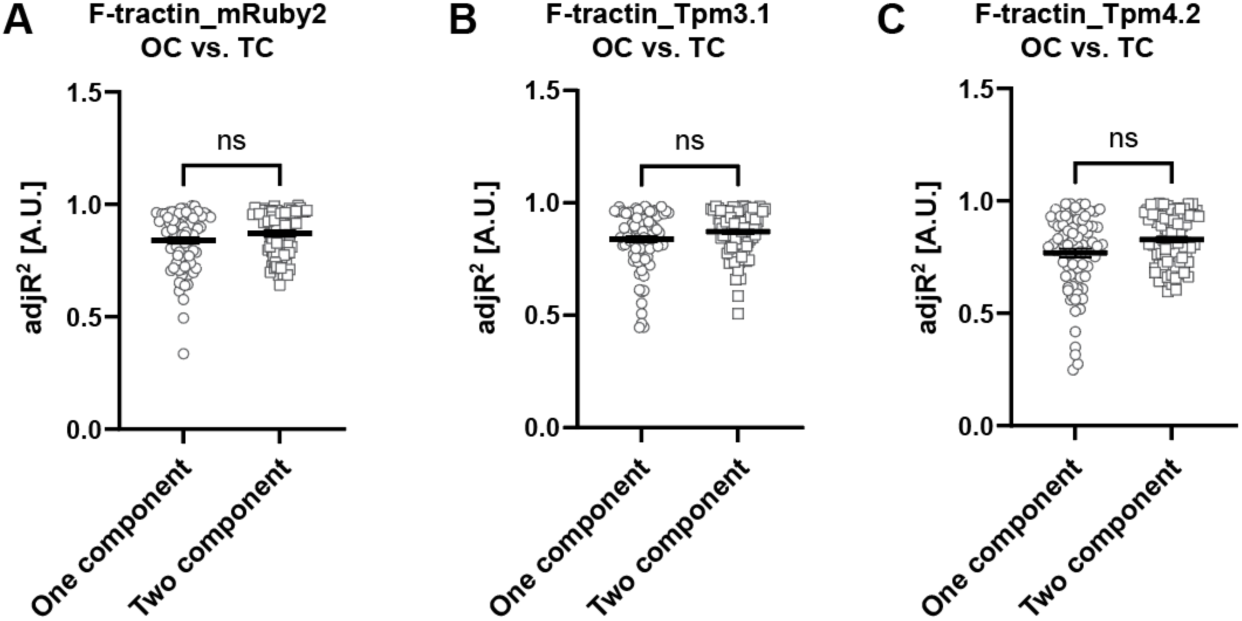
Goodness of the fit (adjR^2^) of OC and TC equations for dendritic spines of primary mouse hippocampal neurons double transduced with F-tractin-EGFP + mRuby2, F-tractin-EGFP + Tpm3.1-IRES-mRuby2 and F-tractin-EGFP + Tpm4.2-IRES-mRuby2 after 120s of recovery. (A) Spines double transduced with F-tractin-EGFP + mRuby2. (B) Spines double transduced with F-tractin-EGFP + Tpm3.1-IRES-mRuby2. (C) Spines double transduced with F-tractin-EGFP + Tpm4.2-IRES-mRuby2. Significance was determined with Mann-Whitney U non-parametric test.

### 2 Supplementary Results

#### 2.1. Dendritic spine fluorescence intensity and surface area for experimental groups mRuby2, Tpm3.1-mRuby2, Tpm4.2-mRuby2

**Figure S3.**
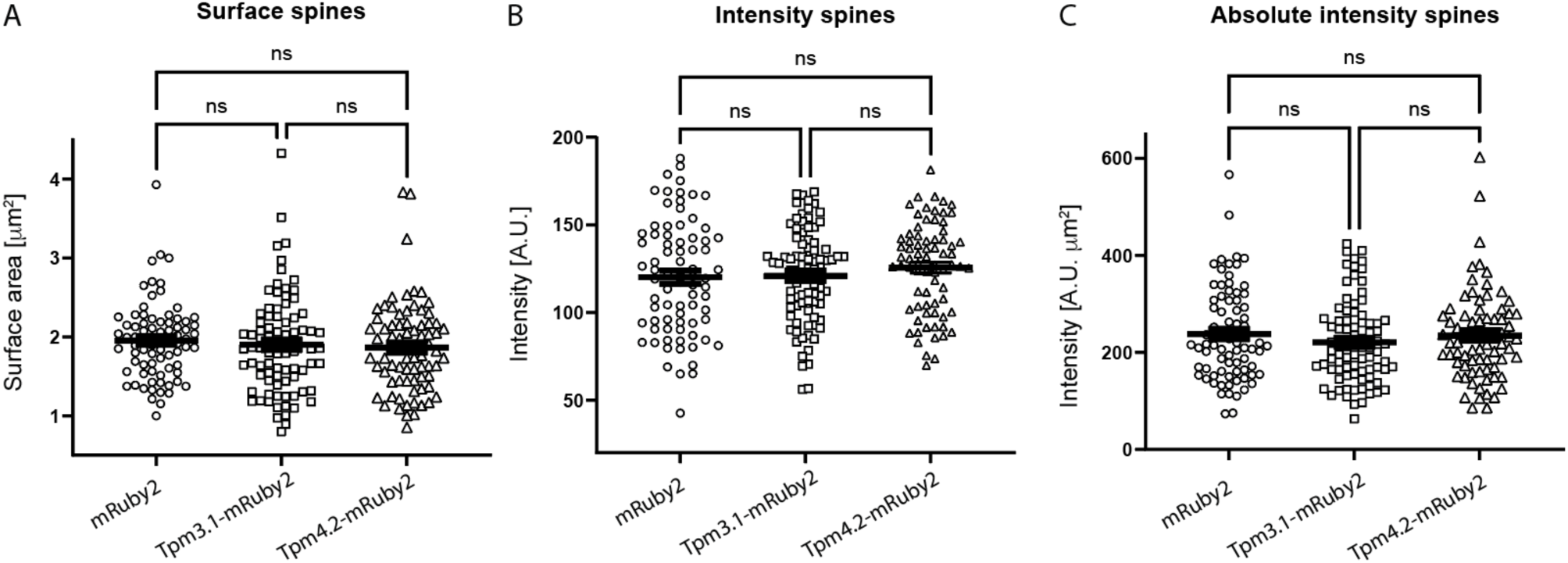
Quantification of mRuby2 fluorophore intensity and the surface of dendritic spines of primary mouse hippocampal neurons transduced with mRuby2, Tpm3.1-mRuby2 or Tpm4.2-mRuby2. (A) Relative mRuby2 intensity in bleached dendritic spines, measured in the first pre-bleaching frame. (B) Absolute mRuby2 intensity in bleached dendritic spines, measured in the first pre-bleaching frame. (C) Surface of bleached dendritic spines, measured in the first pre-bleaching frame. Significance was determined with Mann-Whitney U non-parametric test.

#### 2.2. Dendritic spine fluorescence intensity and surface area for experimental groups F-tractin + mRuby2, F-tractin + Tpm3.1-IRES-mRuby2 and F-tractin + Tpm4.2-IRES-mRuby2

**Figure S4.**
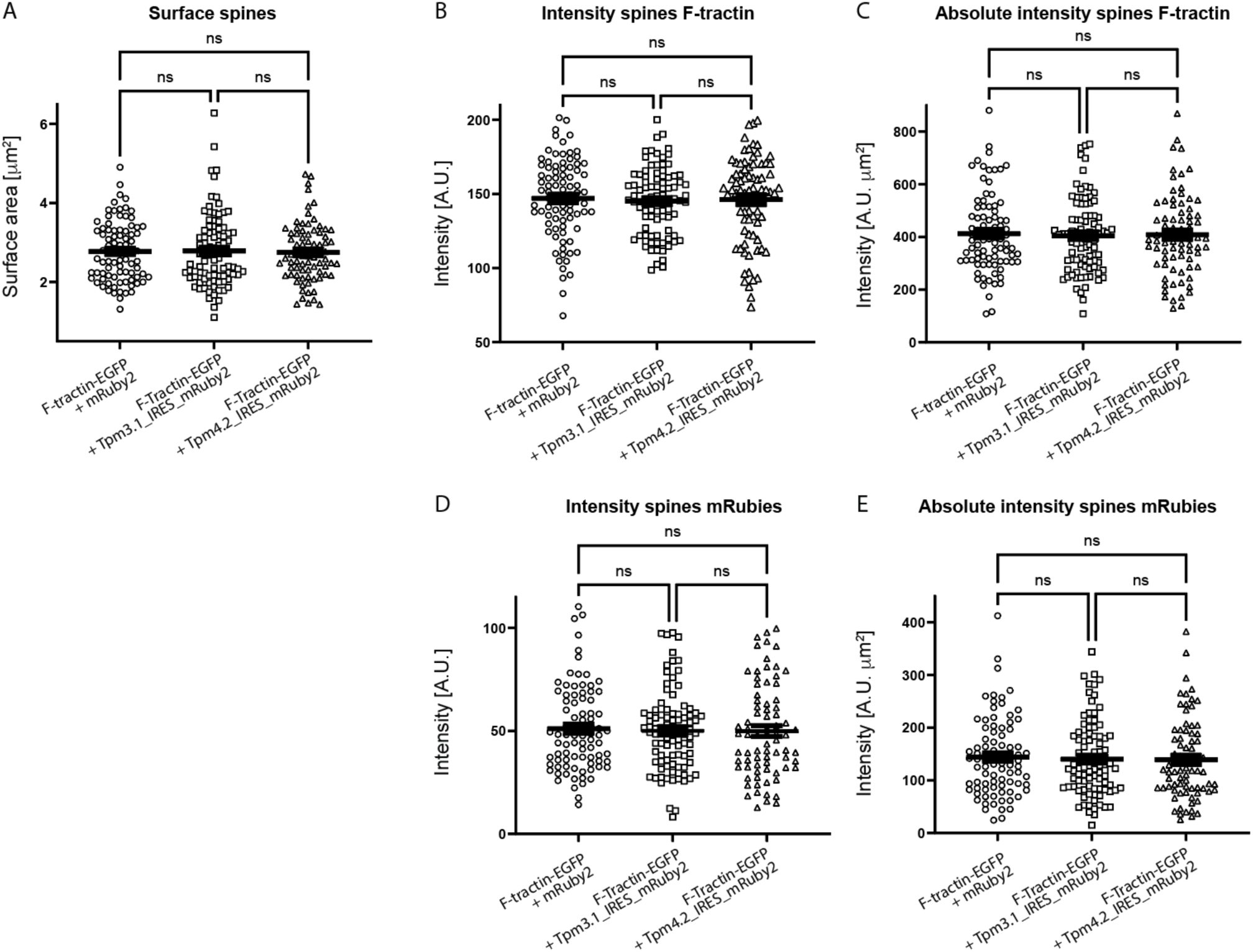
Quantification of EGFP, mRuby2 intensity and surface of dendritic spines of primary mouse hippocampal neurons double transduced with F-tractin-EGFP + mRuby2, F-tractin-EGFP + Tpm3.1-IRES-mRuby2 or F-tractin-EGFP + Tpm4.2-IRES-mRuby2. (A) Relative EGFP intensity in bleached dendritic spines, measured in the first pre-bleaching frame. (B) Relative mRuby2 intensity in bleached dendritic spines, measured in the first pre-bleaching frame. (C) Surface of bleached dendritic spines, measured in the first pre-bleaching frame of merged mRuby2 and EGFP channels. (D)Absolute EGFP intensity in bleached dendritic spines, measured in the first pre-bleaching frame. (E) Absolute EGFP intensity in bleached dendritic spines, measured in the first pre-bleaching frame. Significance was determined with Mann-Whitney U non-parametric test.

## Statements and Declarations

## Author contributions

Conceptualization: TF; Methodology: TT, HS, EP, AC, TF. Formal analysis and investigation: TT and TF; Writing - original draft preparation: TT; Writing - review and editing: TT and TF; Funding acquisition: TF; Resources: TF; Supervision: TF. All authors read and approved the final manuscript.

## Data availability

Additional data will be made available upon request.

## Ethics approval

Approval was obtained from the animal ethics committee of Macquarie University. The procedures used in this study adhere to the tenets of the Declaration of Helsinki.

