## Supplementary figures and images for "Fluorescence recovery after photobleaching reveals different behaviour of tropomyosin isoforms Tpm3.1 and Tpm4.2 in dendritic spines"

### Fig S1.pdf

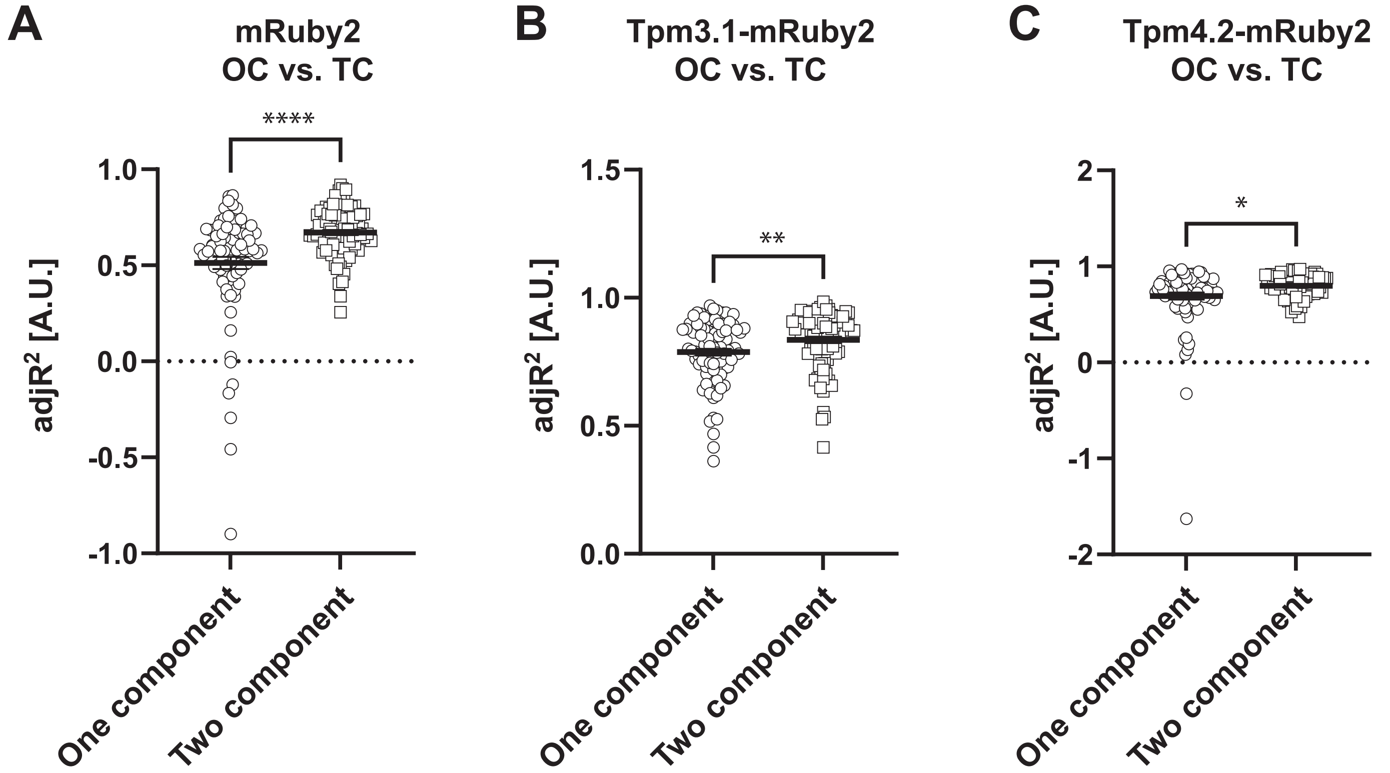

### Fig S2.pdf

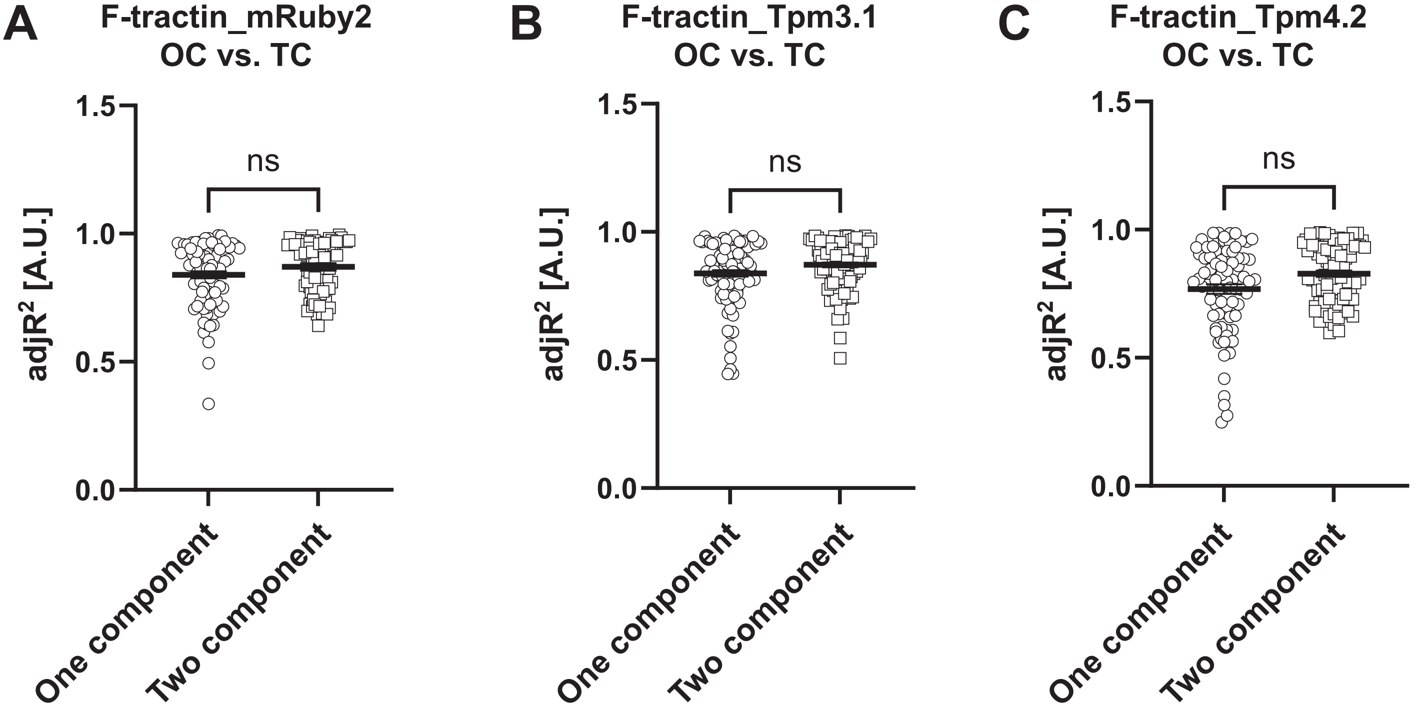

### Fig S3.pdf

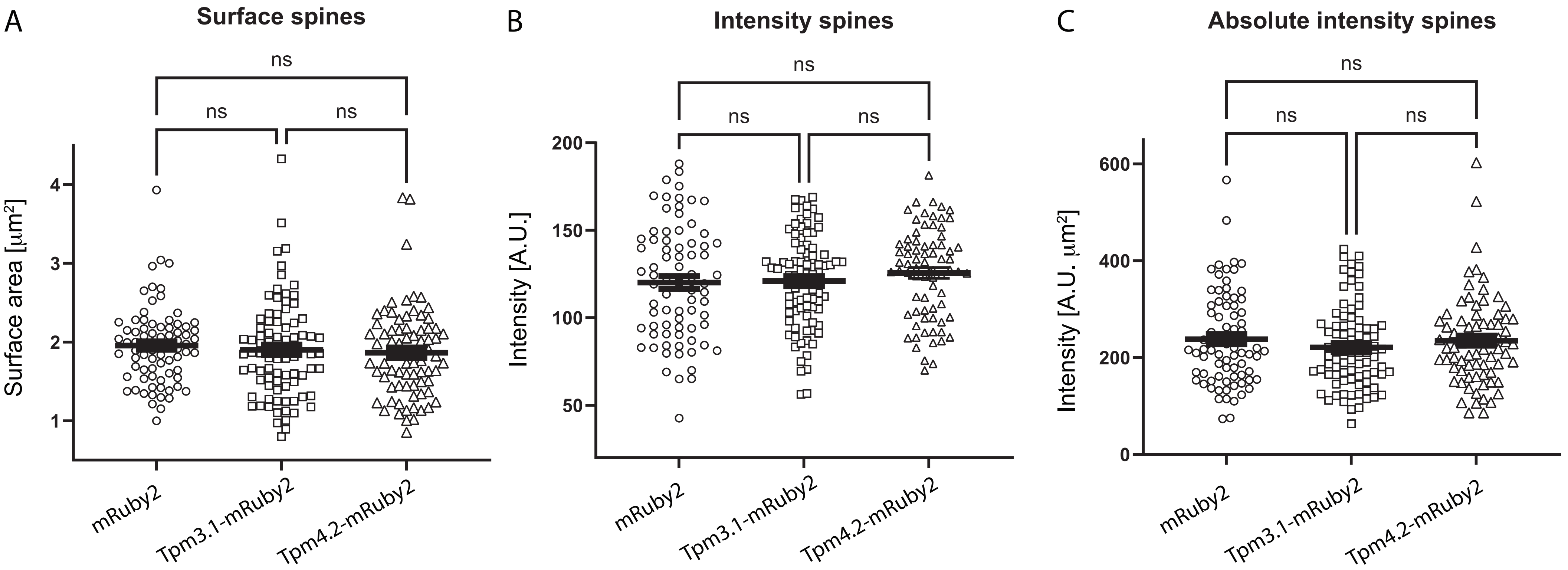

### Fig S4.pdf

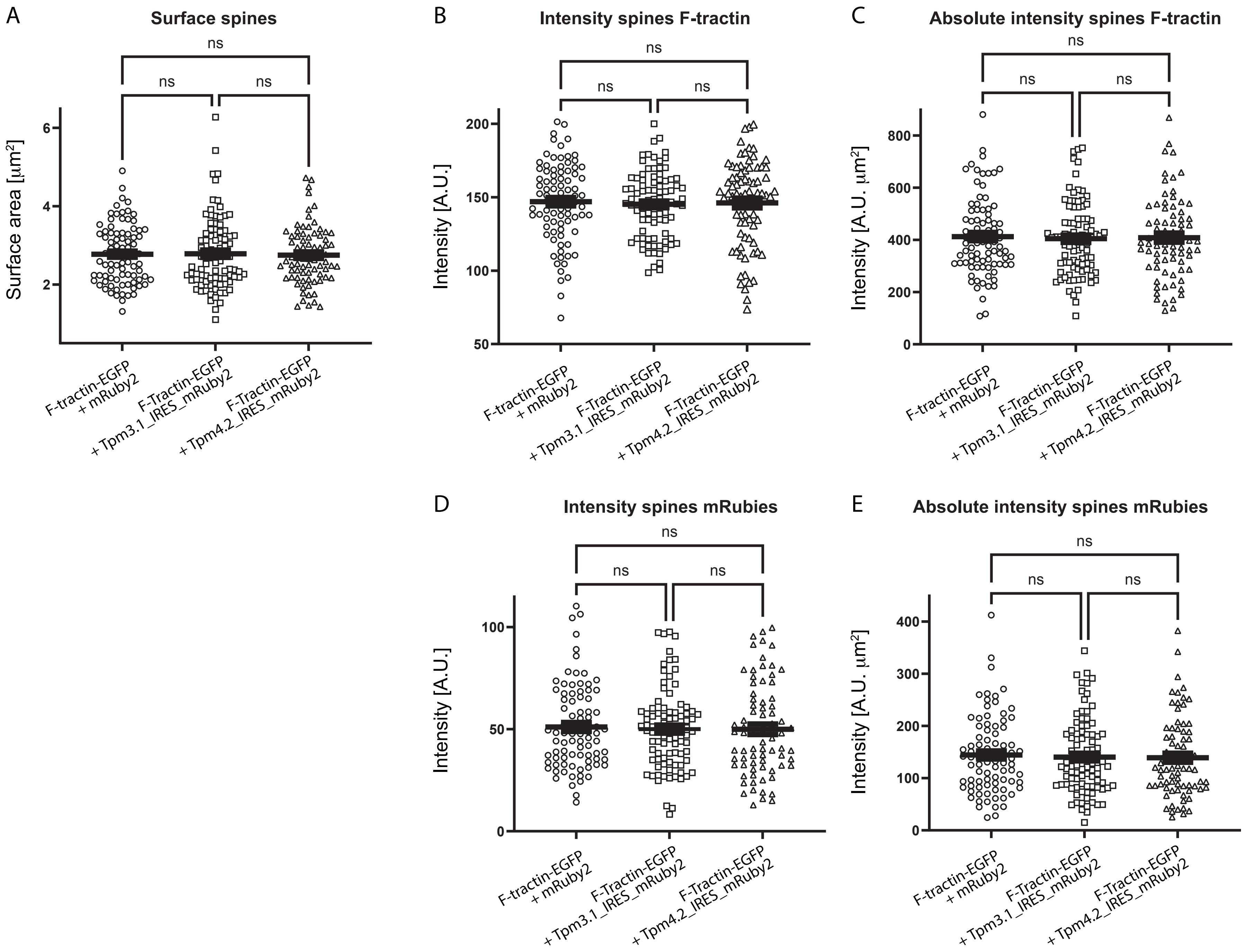
